# Prkra dimer senses double-stranded RNAs to dictate global translation efficiency

**DOI:** 10.1101/2024.07.16.603655

**Authors:** Tong Lu, Pengcheng Ma, Aijun Chen, Hailing Fang, Jianlin Xu, Mingyu Wang, Ling Su, Sen Wang, Yizhuang Zhang, Jiasheng Wang, Boya Yang, De-Li Shi, Yong Zhou, Qianqian Gong, Xiangguo Liu, Bingyu Mao, Ming Shao

**Affiliations:** Shandong Provincial Key Laboratory of Animal Cell and Developmental Biology and Key Laboratory for Experimental Teratology of the Ministry of Education, School of Life Sciences, Shandong University, Qingdao 266237, China; State Key Laboratory of Genetic Resources and Evolution, Kunming Institute of Zoology, Chinese Academy of Sciences, Kunming 650223, China; State Key Laboratory of Microbial Technology, Institute of Microbial Technology, Qingdao 266237, China; Developmental Biology Laboratory, CNRS-UMR7622, Institut de Biologie Paris-Seine, Sorbonne University, Paris 75005, France; Yangtze River Fisheries Research Institute, Chinese Academy of Fishery Sciences, Wuhan 430223, China; Qilu Hospital (Qingdao), Cheeloo College of Medicine, Shandong University, 758 Hefei Road, Qingdao, Shandong, 266035, China; Shandong University-Yuanchen Joint Biomedical Technology Laboratory, Qingdao, 266237, China

**Keywords:** Double-stranded RNA, Prkra, PACT/RAX, dsRNA sensor, translation initiation, early embryos, embryonic stem cells

## Abstract

Double-stranded RNAs (dsRNAs), known as conserved pathogen-associated molecular patterns, are recognized by interferon-induced protein kinase R (PKR) to trigger an integrated stress response, characterized by the inhibition of global translation. However, the interferon system is inactive in pluripotent cells, thus the mechanism underlying dsRNA sensing and translational control is unclear. In this study, we utilized early zebrafish embryos as a model of pluripotent cells and discovered a PKR-independent blockage of translation initiation induced by dsRNA stimulation. Prkra dimer was identified as the genuine dsRNA sensor. Upon binding to dsRNAs, the dimerized dsRNA binding domain 3 becomes activated to sequester the eIF2 complexes from the translation machinery, leading to a hindrance in global protein synthesis. This distinctive embryonic stress response restricts RNA virus SVCV replication in zebrafish embryos but is also conserved in early mouse embryos and embryonic stem cells. Therefore, the Prkra-mediated dsRNA sensing and translation blocking mechanism potentially represents a common strategy for reestablishing physiological homeostasis in response to environmental stresses.

## Introduction

Double-stranded RNAs (dsRNAs) are evolutionarily conserved pathogen-associated molecular patterns that frequently accumulate during viral infections due to viral genome replication.^1–3^ They act as stress stimuli that trigger an integrated stress response (ISR) characterized by a global inhibition of translation.^4–6^ This response is initiated by the activation of the dsRNA sensor protein kinase R (PKR), also known as EIF2AK2.^7^ Mechanistically, upon stimulation by dsRNA, PKR phosphorylates the Ser51 of eIF2α, a subunit of translation initiation factor eIF2. This phosphorylation transforms eIF2 from a substrate into an inhibitor of the guanine nucleotide exchange factor eIF2B.^8,9^ As a result, the conversion of inactive eIF2-GDP to its active form, eIF2-GTP, is impeded. Consequently, the formation of the eIF2 ternary complex and the 43S preinitiation complex, both of which are instrumental in initiating the translation of AUG-started open reading frames, is blocked.^10^ The dsRNA-induced ISR is generally regarded as a part of the antiviral innate immune mechanism that directly inhibits viral protein synthesis.^11–13^ However, since these insights and concepts have been established primarily in differentiated cells, their full applicability to pluripotent stem cells remains unknown.

Accumulating evidence indicates that there are significant differences in the response of undifferentiated and terminally differentiated cells to virus and dsRNA stimulation. In contrast to most cell types, pluripotent embryonic stem cells (ESCs) and embryonic carcinoma cells do not produce type I interferons (IFNs) in response to viral infection or poly(I:C) treatment.^14^ In addition, these cells show weak responses to exogenous IFNs,^15^ suggesting that they do not primarily rely on canonical IFN signaling for antiviral activity. Instead, a panel of intrinsically expressed IFN Stimulated Genes (ISGs) safeguard stem cells against viral infections, with PKR being notably absent from this panel,^16^ which raises questions regarding its involvement in translational control at the early stage of development. Hence, it is plausible that undifferentiated cells employ unique mechanisms for dsRNA recognition and translation regulation, which may not rely on the PKR-eIF2α axis typically observed in differentiated cells.

Interferon-inducible double-stranded RNA-dependent protein kinase activator A (Prkra, also known as PACT in humans, RAX in mice, and Xlrbpa in Xenopus)^17–19^ plays multifaceted roles in cellular stress and beyond. It was originally identified as a protein activator of PKR, modulating translation efficiency and inducing cell death under stress conditions.^17,18,20^ Beyond its role in PKR activation, Prkra is implicated in RNA interference through direct binding to Dicer and can interact with RIG-I to promote type I IFN expression.^21–25^ The antiviral activity of Prkra is underscored by the evolution of several viral proteins that interact with and antagonize its function.^26–28^ Structurally, Prkra contains three tandem dsRNA-binding domains (dsRBDs). The first two domains are responsible for dsRNA binding, while the third domain facilitates dimerization and protein-protein interactions.^17,29–33^ As aforementioned, RIG-I-mediated type I IFN signaling and PKR are likely nonfunctional in early embryos and stem cells. This phenomenon raises a pertinent question as to whether Prkra possesses uncharacterized functions independent of PKR, RNA interference and IFN signaling during early development.

Using zebrafish early embryos as a model of pluripotent cells, we discovered that dsRNAs trigger a distinct stress response characterized by the inhibition of translation. Contrary to the current understanding, the classic players PKR and phosphorylated eIF2α do not contribute to this translation repression. Instead, Prkra functions as a conserved dsRNA sensor controlling global translation rate in early zebrafish embryos and mouse ESCs (mESCs). Distinct from eIF2α phosphorylation, Prkra forms dimers and mediates global inhibition of translation by directly sequestering the eIF2 complex from the translation machinery. Therefore, our work has uncovered a conserved mechanism of global translational control that is initiated by a novel dsRNA sensor.

## Results

### The dsRNAs induce an ISR-like translation blockage in early zebrafish embryos

Early zebrafish embryos serve as an ideal model for studying the effects of dsRNA stimulation on pluripotent cells, owing to the ease of microinjection and the availability of large numbers of embryos. Injection of dsRNAs into early zebrafish embryos induced unspecific toxic effects.^34^ Embryos injected with dsRNA either lysed during the gastrula stage or exhibited lethal malformations by 36 hours post-fertilization (hpf). These aberrant phenotypes include tail truncation, split axis, abnormalities or absence of eyes, malformed brain structures, and cardiac edema (Figure S1A). Injection of dsRNAs (100 pg/embryo) with different lengths (285 bp-1043 bp) and sequences (Figure S1B and Table S1) induced similar lethal phenotypes (Figures S1C and S1D). These results suggest that dsRNA, as a molecular pattern, may trigger a robust stress response in the early embryo in a sequence-independent manner. We thus mainly used a 1 kb dsRNA (dsRNA2 in Figure S1B) with sequence from *cas9* open reading frame in the following experiments.

There is a possibility that the toxic effects arise due to a widespread translation blockage by dsRNA-induced ISR. To verify this hypothesis, we injected *gfp* reporter mRNAs along with dsRNAs at different doses. Compared with injection of *gfp* mRNAs alone, coinjection of dsRNAs (30 pg/embryo) with *gfp* mRNAs remarkably reduced GFP protein levels by over 90% (Figure 1A and B), and coinjection of dsRNAs at a lower dose (10 pg/embryo) with *gfp* mRNAs also significantly reduced GFP protein synthesis, albeit to a lesser extent (Figures 1B). We also tested luciferase reporter mRNAs in the coinjection experiment and obtained similar results (Figure 1C). There must be a translation blockage because the integrity and expression levels of *gfp* mRNAs were unaffected after dsRNA coinjection (30 pg/embryo), as determined by northern blot analysis (Figure 2D). Next, we performed puromycin incorporation experiments to determine if global translation inhibition occurs upon dsRNA stimulation. Similar to the positive control in which cycloheximide (CHX) treatment was used to block translation, we observed a dramatic decrease in puromycin-labeled protein levels after injection of dsRNAs at 30 or 100 pg/embryo (Figures 1E-1G). By analyzing the ribosome profile through sucrose density gradient ultracentrifugation, it was evident that dsRNA stimulation significantly reduced the levels of polysomes (Figures 2H and I). These data collectively confirmed the occurrence of a global translation repression in response to dsRNA stimulation in early zebrafish embryos.

**Figure 1.**
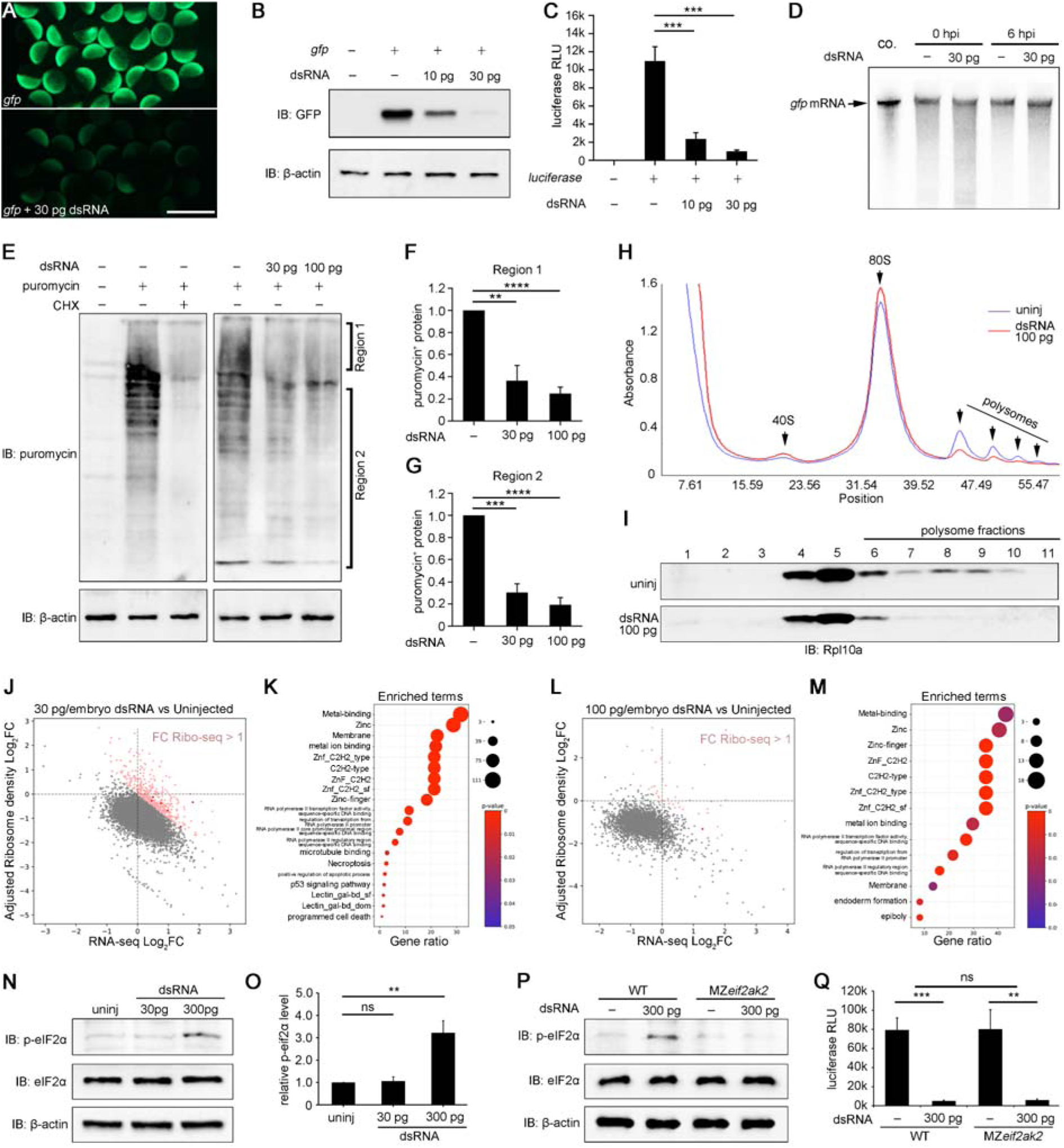
dsRNAs induce an ISR-like global translation repression. (**A**) Significant suppression of GFP expression in embryos injected with *gfp* mRNA and dsRNAs (30 pg/embryo), as revealed by fluorescent imaging. (**B** and **C**) Representative western blot results and quantification. (**D**) Northern blot analysis shows the absence of degradation of *gfp* reporter mRNAs by dsRNAs. (**E**) Puromycin incorporation shows significantly reduced newly synthesized proteins after injection of dsRNAs in zebrafish embryos. (**F** and **G**) Quantification of regions 1 and 2. (**H**) Ribosome profile comparison between uninjected and dsRNA-injected embryos at 6 hpf. (**I**) Western blot analysis of ribosome large subunit marker Rpl10a in fractions separated by sucrose gradient ultracentrifugation. (**J** and **L**) Changes in mRNA abundance and translation efficiency (adjusted ribosome density) after stimulation with low dose (**J**) and high dose (**L**) dsRNAs. Pink-red designates genes with enhanced protein synthesis rates. (**K** and **M**) Enriched terms of genes with upregulated protein synthesis. (**N** and **O**) Phosphorylation of eIF2α induced by high-dose of dsRNAs (300 pg/embryo). (**P**) Failure to phosphorylate eIF2α in MZ*eif2ak2* embryos by dsRNAs (300 pg/embryo). (**Q**) Comparable dsRNA-induced translation blockage between wild-type (WT) and MZ*eif2ak2* embryos. Values are means ± SDs. ** p<0.01, *** p<0.001, **** p<0.0001, ns, p>0.05, Student’s *t*-test. ns in **Q**, p>0.05, two-way ANOVA. Scale bar: 1 mm

**Figure 2.**
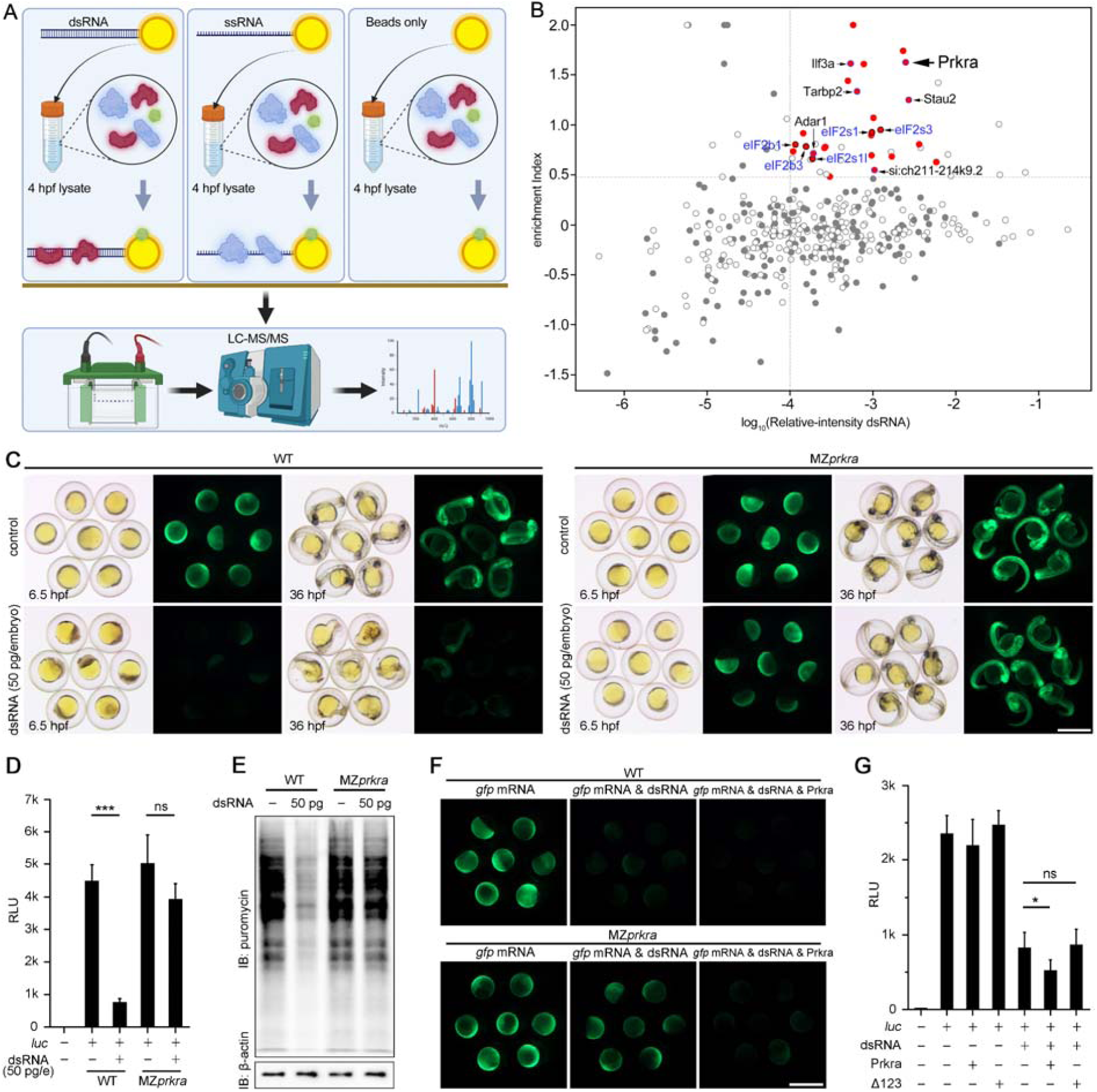
Identification of Prkra as dsRNA sensor. (**A**) Experimental design of RNA-pulldown followed by MS. (**B**) Scatter plot showing enrichment with respect to relative intensity. Specific dsRNA-associated components are highlighted by red dots (see Materials and Methods and Data S3). Known dsRBPs are in black; eIF2 and eIF2B components are in blue. Grey circles represent proteins with high contamination in RNA-free beads. (**C**) Comparison of *in vitro* transcribed *gfp* mRNA translation in wild-type embryos and MZ*prkra* mutants in response to dsRNA stimulation. (**D**) Analysis of *luciferase* mRNA translation in wild-type embryos and MZ*prkra* mutants in the presence of dsRNAs. (**E**) Puromycin incorporation showing decreased global protein synthesis upon dsRNA stimulation. (**F**) Prkra enhances translation blockage by dsRNAs (30 pg/embryo) in both wild-type embryos and MZ*prkra* mutants. (**G**) Translation blockage by dsRNAs in the rabbit reticulocyte *in vitro* translation system is enhanced by recombinant Prkra but not mutant Prkra (with the lengthy α-helix in each dsRBD deleted: Δ123). Values shown are means ± SDs. * p<0.05, *** p<0.001, ns p>0.05, Student’s *t*-test. Scale bars: 1 mm.

To further characterize the dsRNA-induced global translation blockage, we performed ribosome profiling (Ribo-seq) analysis of dsRNA-injected embryos at 6 hpf,^35^ after injection of either low dose (30 pg/embryo) or high dose (100 pg/embryo) of dsRNAs. Ribo-seq has collectively generated 16.07 M uniquely mapped reads. The high quality of Ribo-seq data was confirmed by the length of ribosome-protected fragments (RPFs), the triplet periodicity of 5’ ends mapping of RPFs, and the pairwise correlation of two replicates (Figures S2A-S2E). The mRNAs exhibited a broad range (200-500 folds) of translation efficiency (Figures 1J and L). After adjusting for global translation inhibition as determined by puromycin incorporation (see Figure S2F and Materials and Methods), more than 97.2% of transcripts exhibited decreased translation efficiency following dsRNA injection (Figures 1J and 1L, Data S1). Protein production after dsRNA stimulation was also estimated by assaying the adjusted RPF counts of each gene. The protein synthesis rate of 389 genes, representing 4.41% of tested genes, remained unaffected or even enhanced after injecting 30 pg/embryo of dsRNAs (Figure 1J and Data S1). GO analysis highlighted terms related to zinc finger and sequence-specific DNA binding (Figure 1K). However, only 43 genes, or 1.25% of tested genes, showed a high rate of protein synthesis when 100 pg/embryo of dsRNAs were injected (Figure 1L and Data S1). GO analysis revealed a similar enrichment of zinc finger-related transcription factors (Figure 1M). Surprisingly, in the background of the robust global translation blockage, dsRNA stimulation did not alter the normalized occupancy of ribosomes either at the 5’ regions of open reading frames (ORFs) or the positions near the stop codon (Figures S2G and S2H). As Ribo-seq only captures the translation process after loading the ribosome onto mRNAs, the similar distribution of RPFs suggests that dsRNAs may primarily affect the early phase of translation initiation. However, translation elongation appears to proceed normally. These findings demonstrate an ISR-like translatomic reshaping following injection of dsRNAs into early embryos.

### PKR (Eif2ak2) and phospho-eIF2**α** are dispensable for dsRNA-induced translation inhibition in early embryos

Given that the PKR-eIF2α axis mediates repression of translation initiation,^8,9^ we sought to determine its role in pluripotent cells. Intriguingly, *eif2ak2* was expressed in trace amounts in the early zebrafish embryo (Data S2).^36^ Injecting dsRNAs at a dose of 30 pg/embryo led to profound translation inhibition, but did not trigger the phosphorylation of eIF2α, which was only observed at an extremely higher dose (300 pg/embryo) of dsRNAs (Figure 1N and O). These results suggest that the PKR/eIF2α axis is unlikely to be responsible for dsRNA-induced translation inhibition.

To definitively address this issue, we generated maternal-zygotic mutants for *eif2ak2* (MZ*eif2ak2*) through the “CRISPant” approach.^37^ Genotyping confirmed the absence of wild-type cDNA in the offspring (Figure S3A). MZ*eif2ak2* embryos developed normally and are viable (Figure S3B). In contrast to wild-type embryos, MZ*eif2ak2* embryos injected with dsRNA at 300 pg/embryo did not exhibit eIF2α phosphorylation (Figure 1P), indicating a disruption in residual PKR activity. However, translation remained robustly inhibited in the mutants, comparable to wild-type embryos (Figure 1Q). These findings strongly suggest that dsRNAs induce ISR-like phenomena in early embryos through a distinct dsRNA sensor that operates independently of the PKR and phospho-eIF2α.

### Prkra is the dsRNA sensor mediating translation blockage in the early embryo

We next performed stringent RNA-pulldown followed by mass spectrometry (MS) experiments using biotinylated dsRNAs (120 bp, with 5’-end biotinylated; sequence shown in Table S1) as bait, to identify the genuine dsRNA sensor responsible for translation control in zebrafish early embryos. Single-stranded RNAs (ssRNAs, the biotinylated strand of the dsRNAs) and RNA-free beads were used as negative controls (Figure 2A). By analyzing the abundance and specificity of proteins pulled down by dsRNAs (see Methods), we identified a total of 26 proteins (Figure 2B and Data S3), including six known dsRNA-binding proteins (dsRBPs) and five subunits of the eIF2 and eIF2B complexes. In particular, GO analysis highlighted “dsRNA binding”, “translation initiation”, “eIF2 complex” and “eIF2B complex” in the enriched terms (Figure S4).

Among the potential dsRBPs, Prkra stands out as a compelling candidate for a dsRNA sensor. It demonstrated an exceptional enrichment of over 70-fold in pulldown experiments using dsRNAs compared to ssRNAs and exhibited one of the highest signal intensities (Figure 2B and Data S3). For functional validation, we next generated MZ*prkra* mutants using the CRISPR/Cas9 approach. Disruption of the *prkra* gene resulted in nonsense-mediated decay occurring in all F1 early embryos, as demonstrated by the results of in situ hybridization (ISH) and RT-PCR (Figures S5A and S5B), indicating complete removal of *prkra* gene products.

Prkra mutation did not affect embryonic development (Figure S5C). However, injection of 50 pg dsRNAs into the MZ*prkra* embryo resulted in very mild or no phenotypes, with most embryos developing normally. In contrast, an equal amount of dsRNAs in wild-type embryos induced lethal malformations (Figures S5D and S5E). Consistent with the remarkably weakened phenotypes, *prkra* mutation significantly alleviated the translational inhibition of *in vitro* transcribed *gfp* and *luciferase* mRNAs (150 pg/embryo each) mediated by dsRNAs (Figures 2C and D). The dsRNA-mediated blockage of puromycin incorporation was also released in MZ*prkra* mutant embryos (Figure 2E). Conversely, the co-injection of recombinant Prkra protein into wild-type embryos enhanced translation inhibition, whereas co-injection into MZ*prkra* embryos rescued the defect in response to dsRNA stimulation, allowing for subsequent activation of translation suppression (Figure 2F). Significantly, in a reticulocyte lysate *in vitro* translation system, the addition of dsRNAs led to a decreased translation, and the presence of wild-type but not mutant Prkra (Δ123, with the lengthy α-helix in each dsRBD deleted) further enhanced the translational repression (Figure 2G). These results collectively indicate that Prkra acts as a dsRNA sensor, playing a crucial role in global translational control.

### Prkra binds to dsRNAs as dimers in a highly organized manner

Next, we investigated the mechanisms by which Prkra regulates global translation. We first conducted biochemical analyses to validate its dsRNA-binding capacity and specificity. Consistent with the MS data, recombinant Prkra protein was specifically pulled down by dsRNAs, but not by ssRNAs (Figure 3A). Conversely, when using anti-His beads to precipitate Prkra-6xHis, only dsRNAs were effectively co-precipitated (Figure 3B). The electrophoretic mobility shift assay (EMSA) experiments also demonstrated the robust binding capacity of Prkra to dsRNAs (Figure 3C). The molecular weight of the Prkra-dsRNA complexes was significantly increased with the addition of more proteins, indicating the formation of dsRNA-associated Prkra complexes (Figure 3C). Unlabeled dsRNAs were able to compete with biotinylated dsRNAs for binding to Prkra, as evidenced by the decreased molecular weight of the Prkra-dsRNA complexes (Figure 3C).

**Figure 3.**
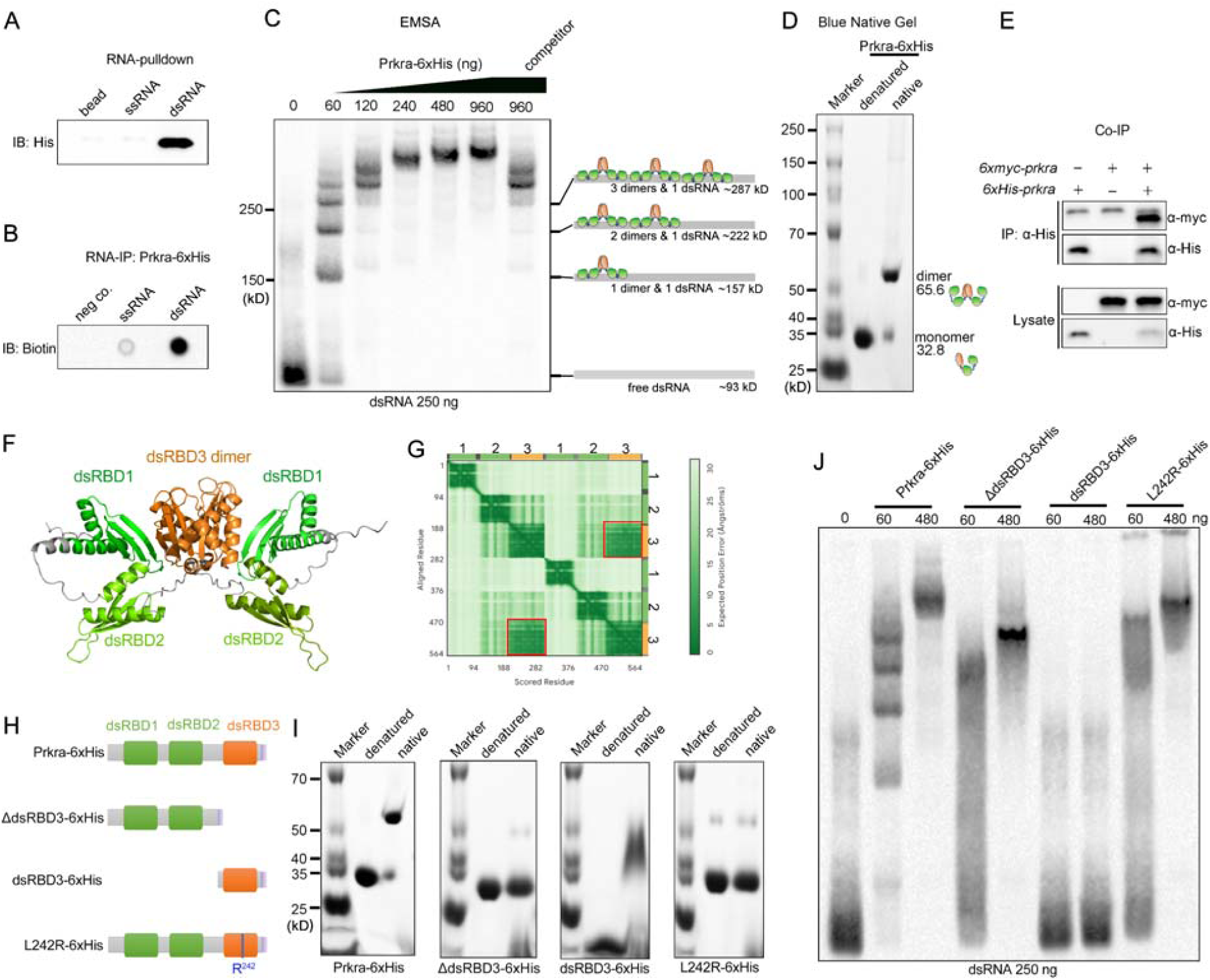
Dimerization of Prkra is essential for highly ordered binding of dsRNAs. (**A**) RNA pulldown shows the specific interaction of Prkra-6xHis with dsRNAs. (**B**) RNA IP shows the specific binding of biotin-labeled dsRNAs by His-tagged Prkra. (**C**) EMSA analysis reveals a laddering pattern of Prkra-dsRNA complexes with an approximate 65 kDa interval between adjacent bands, indicating the dimerization of Prkra when binding to dsRNAs. The competitor means addition of 2.5 μg unlabeled dsRNAs. (**D**) Dimer formation of Prkra-6xHis revealed by native blue gel analysis. (**E**) The overexpression of 6xMyc-Prkra and 6xHis-Prkra in zebrafish embryos leads to the formation of complexes, as demonstrated by co-IP analysis. (**F**) Predicted putative structure of Prkra by Alphafold2. Orange highlights dimerized dsRBD3. (**G**) The expected positional error plot of the predicted structure highlights dimer signals between the dsRBD3, as indicated by the red boxes. (**H**) Schematic representation illustrating wild-type Prkra and its mutant forms. (**I**) Dimerization assays of mutant Prkra forms using native blue gel electrophoresis demonstrate that wild-type Prkra and dsRBD3, but not the L242R mutant and ΔdsRBD3, are capable of forming dimers. (**J**) EMSA analysis shows the absence of interaction between dsRBD3 and dsRNAs. Note that the dimerization-defective mutants L242R and ΔdsRBD3 do not exhibit the laddering pattern characteristic of wild-type Prkra when binding to dsRNAs.

Surprisingly, we observed that Prkra-dsRNA complexes appear as ladders. By incorporating molecular weight markers in EMSA, we determined that there is an approximate 65 kD interval between each two bands. As the molecular weight of the Prkra-6xHis protein is 32.8 kD, the presence of a 65 kD interval in the laddering pattern suggests that Prkra-6xHis may bind to dsRNAs as dimers (Figure 3C). Further analysis using blue native gel electrophoresis revealed that native Prkra-6xHis exhibited a molecular weight ranging from 50-70 kD, approximately twice the size of the denatured Prkra monomer, which displayed a molecular weight slightly lower than 35 kD (Figure 3D). To further substantiate these findings, we injected zebrafish embryos with *6xmyc-prkra* and *6xHis-prkra* mRNAs and conducted co-immunoprecipitation (co-IP) analysis with anti-His antibody (Figure 3E). The results demonstrated that 6xmyc-tagged Prkra co-immunoprecipitated with 6xHis-Prkra, further confirming dimeric status of Prkra in the embryos.

Interestingly, AlphaFold3 predicts dimer formation through dsRBD3 (Figure 3F), as evidenced by the low Expected Position Error observed between the two dsRBD3 domains in the Predicted Aligned Error (PAE) plot (Figure 3G).^38^ This prediction is consistent with previous studies demonstrating that human PRKRA/PACT can form dimers via dsRBD3 *in vitro*. The L263R mutation in human PRKRA/PACT disrupts dimerization.^33^ To investigate the impact of this single amino acid substitution on dimerization of zebrafish Prkra, we generated an analogous L242R mutant of His-tagged zebrafish Prkra protein. Additionally, we created a ΔdsRBD3-6xHis mutant, which lacks the dsRBD3 domain of Prkra, alongside the intact dsRBD3-6xHis as controls (Figure 3H). Native blue gel analysis demonstrated that both the wild-type Prkra and the dsRBD3 were capable of forming dimers. In contrast, neither the L242R mutant nor the ΔdsRBD3 were able to dimerize (Figure 3I).

Next, we utilized these Prkra mutants to investigate the impact of dimerization on their binding affinity for dsRNAs. EMSA analysis revealed that dsRBD3-6xHis alone does not bind to dsRNAs (Figures 3J and S6); however, binding to dsRNAs remains intact following the deletion or mutation of dsRBD3 (Figure 3J and S6). Surprisingly, the characteristic laddering pattern was lost upon the addition of the L242R-6xHis and ΔdsRBD3-6xHis mutants, which instead exhibited a smeared binding pattern in the EMSA analysis (Figure 3J and S6). This suggests a disordered interaction between these dimerization-defective Prkra proteins and dsRNAs. Given that EMSA is performed on a non-denaturing gel, the formation of distinct bands necessitates the presence of well-ordered complexes with nearly identical electrophoretic mobilities. Only complexes with similar folding conformations display consistent migration rates. These results indicate that while dimerization is not essential for Prkra binding to dsRNAs, which is primarily mediated by dsRBD1 and dsRBD2, it is crucial for the formation of a highly organized binding structure between Prkra and dsRNA.

### Prkra is activated by dsRNAs and uses its dimerized dsRBD3 to grasp the eIF2 complex

The RNA pulldown and MS data revealed a specific interaction of dsRNAs with the eIF2 and eIF2B complexes (Figures 2B). This suggests that the dsRNA-Prkra complex may sequester the eIF2 complex, effectively pulling it away from the translation machinery. To test this hypothesis, we first confirmed the specific interaction between dsRNAs and eIF2α both *in vitro* and *in vivo* through RNA pulldown and western blot analyses, demonstrating that dsRNAs, unlike ssRNA, could effectively pull down eIF2α (Figures 4A and 4B). Importantly, this interaction was dependent on the presence of Prkra, as shown by the significantly reduced binding of eIF2α to dsRNAs when using lysates from MZ*prkra* embryos at 4 hpf (Figure 4C). Conversely, the addition of recombinant Prkra-6xHis to lysates derived from either wild-type or MZ*prkra* embryos at 4 hpf resulted in a substantial increase in the amount of pulled-down eIF2α (Figure 4C).

**Figure 4.**
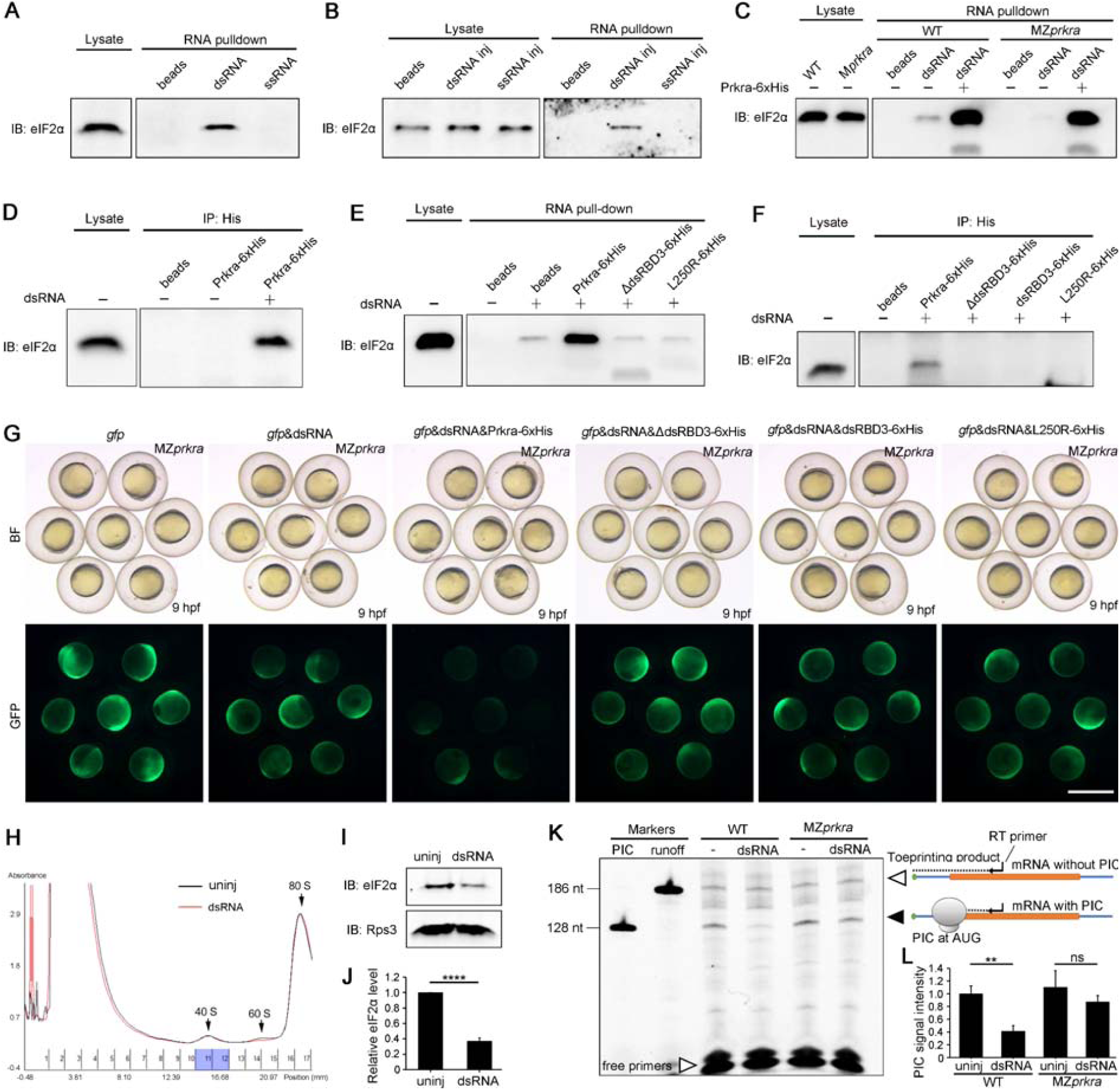
Prkra dimer is activated by dsRNAs to sequester eIF2 from preinitiation complex. (**A**) Biotinylated dsRNAs, but not ssRNAs, effectively pull down eIF2α from the lysate of wild-type zebrafish embryos at 4 hpf. (**B**) Injected biotinylated dsRNAs interact with eIF2α *in vivo*, as demonstrated by co-purification with dsRNA, whereas biotinylated ssRNAs do not show interaction with eIF2α. (**C**) The absence of Prkra diminished the binding of dsRNAs to eIF2α, whereas the addition of Prkra-6xHis significantly enhanced the sequestration of eIF2α. (**D**) The presence of dsRNAs is essential for Prkra to interact with eIF2α, as demonstrated in Co-IP analysis. (**E**) Dimerization-defective Prkra mutants ΔdsRBD3 and L242R are unable to enhance the capacity of dsRNAs in pulling down eIF2α. (**F**) Co-IP analyses demonstrate the requirement of Prkra to interact with eIF2α in the presence of dsRNA, whereas dsRBD3 dimer alone is insufficient to bind to eIF2α. (**G**) Wild-type Prkra, but not dsRBD3 or the dimerization-defective mutant Prkra, induces translation repression in MZ*prkra* mutant embryos stimulated by dsRNAs. (**H**) Absorbance at 260 nm of sucrose gradient ultracentrifugation of samples from embryos at 6 hpf with or without dsRNA injection (100 pg/embryo). Fractions containing the 43S preinitiation complex is highlighted by blue. (**I** and **J**) Significant decrease in 43S preinitiation complex (PIC) in the presence of dsRNAs, with eIF2α serving as a PIC marker. (**K**) Toeprinting analysis shows a significant reduction in PIC formation on the start codon of *gfp* mRNAs in the lysates of dsRNA-injected (100 pg/embryo) wild-type embryos, but not MZ*prkra* embryos. (**L**) Quantification of PIC assembly on *gfp* mRNA in different lysates as indicated. Values shown are means ± SDs. ** p<0.01, **** p<0.0001, ns p>0.05, Student’s *t*-test. Scale bars: 1 mm.

We next investigated whether the interaction between Prkra and the eIF2 complex is dependent on dsRNAs. Recombinant Prkra-6xHis was incubated with lysates from wild-type embryos at 4 hpf, both in the presence and absence of dsRNA. The Prkra-associated complexes were then precipitated using an anti-His antibody. Notably, no eIF2α was co-purified with Prkra-6xHis in the absence of dsRNAs (Figure 4D). In sharp contrast, the presence of dsRNAs significantly enhanced the sequestration of eIF2α by Prkra (Figure 4D), indicating that dsRNAs activate Prkra to engage with the eIF2 complex.

As the dimerization of Prkra dictates its highly organized binding to dsRNAs, it is thus intriguing to investigate whether this phenomenon influences the dsRNA-dependent interaction between Prkra and the eIF2 complex. Notably, the addition of Prkra-6xHis to the lysate from wild-type embryos at 4 hpf significantly enhanced the amount of eIF2α pulled down by dsRNAs (Figure 4E). In contrast, this effect was not observed when using dimerization-defective mutant proteins, ΔdsRBD3-6xHis and L242R-6xHis (Figure 4E). We also conducted co-immunoprecipitation (co-IP) experiments to validate these findings. As anticipated, in the presence of dsRNAs, the ΔdsRBD3-6xHis and L242R-6xHis mutant proteins were unable to bind to eIF2α, whereas wild-type Prkra successfully did so (Figure 4F). Notably, dsRBD3-6xHis, which was capable of dimerization but could not bind dsRNAs, also failed to interact with the eIF2 complex (Figure 4F). Consistent with these observations, the supplementation of dsRBD3-6xHis, ΔdsRBD3-6xHis, or L242R-6xHis failed to recover the inhibition of *gfp* mRNA translation in MZ*prkra* embryos stimulated by dsRNAs, in stark contrast to the effects observed with wild-type Prkra-6xHis (Figure 4G). These results demonstrate that dimerization of dsRBD3 in Prkra is crucial for its interaction with eIF2, and that this interaction is further dependent on the proper structural alignment of the dsRBD3 dimers on dsRNAs.

Considering that eIF2 plays a crucial role in the assembly of the 43S preinitiation complex to regulate the rate of translation initiation, it is critical to determine whether binding of eIF2 to the Prkra-dsRNA complex is sufficient to reduce the formation of the 43S preinitiation complex *in vivo*. To address this question, we conducted fragmentation experiments using sucrose density gradient ultracentrifugation. Fragments located near the 40S ribosome region were isolated based on the absorbance profile for light excited at 260 nm (Figure 4H). Notably, the amount of eIF2α, serving here as a marker for the 43S preinitiation complex, exhibited a significant decrease upon dsRNA stimulation (Figures 4I and J), which corresponded to the observed reduction in global protein synthesis levels.

To directly visualize the formation of the pre-initiation complex at the start codon, we conducted a toeprinting assay using *in vitro* translation systems derived from wild-type and MZ*prkra* embryos at 4 hpf, with or without the injection of dsRNAs. By adding *in vitro* transcribed *gfp* mRNA as a model mRNA, we were able to detect the preinitiation complex at the AUG codon of *gfp* mRNA due to its ability to inhibit primer extension during the reverse transcription reaction. As anticipated, in the toeprinting assay using lysates from wild-type embryos injected with dsRNAs, the pre-initiation complex signal of 128 nt was reduced to 41% of the control level (Figure 4K and L). In contrast, this reduction was not observed in the group using lysates from MZ*prkra* embryos that were also injected with dsRNAs (Figure 4K and L), which is consistent with the observation that dsRNAs fail to induce translation blockage in the absence of Prkra. Collectively, these results identified a dsRNA-activated Prkra dimer-dependent eIF2 complex sequestering mechanism underlying global translation blockage.

### Prkra-mediated translation repression restricts virus replication in early zebrafish embryos

We next aimed to elucidate the biological significance of Prkra in translation remodeling. Given that RNA virus replication is a common source of dsRNAs, it is highly plausible that the anti-viral activity associated with Prkra-mediated translation inhibition plays a crucial role in this context. To address this hypothesis, we introduced spring viremia of carp virus (SVCV), a negative-sense RNA virus, into zebrafish embryos to determine whether the dsRNA-induced response serves as a valid antiviral mechanism in early embryonic development. Embryos lacking Prkra are significantly more vulnerable to SVCV infection, with most of them exhibiting severe malformations by 12 hpf (Figure 5A). As anticipated, these severe phenotypes are associated with uncontrolled viral replication. Quantitative reverse transcription PCR (qRT-PCR) analysis revealed that SVCV displayed an explosive replication trend in MZ*prkra* mutants compared to wild-type embryos (Figure 5B). Strikingly, the intentional injection of dsRNAs (50 pg/embryo) to activate Prkra resulted in a nearly complete inhibition of SVCV replication (Figure 5C). Consistently, six hours after injection of the virus into wild-type embryos, we found a significant inhibition of translation, as demonstrated by the puromycin incorporation assay, which showed an approximately 50% overall reduction of protein levels (Figure 5D). The expression levels of IFN ligands were low (Data S2), and could not be activated following viral infection in zebrafish embryos (Figure 5E), which supports the observation that dsRNAs failed to induce IFN expression (Figure 5F). These results indicate that the translation repression triggered by the dsRNA sensor Prkra constitutes a fully functional antiviral stress response in early zebrafish embryos, thus excluding the involvement of the IFN system in the restriction of SVCV replication.

**Figure 5.**
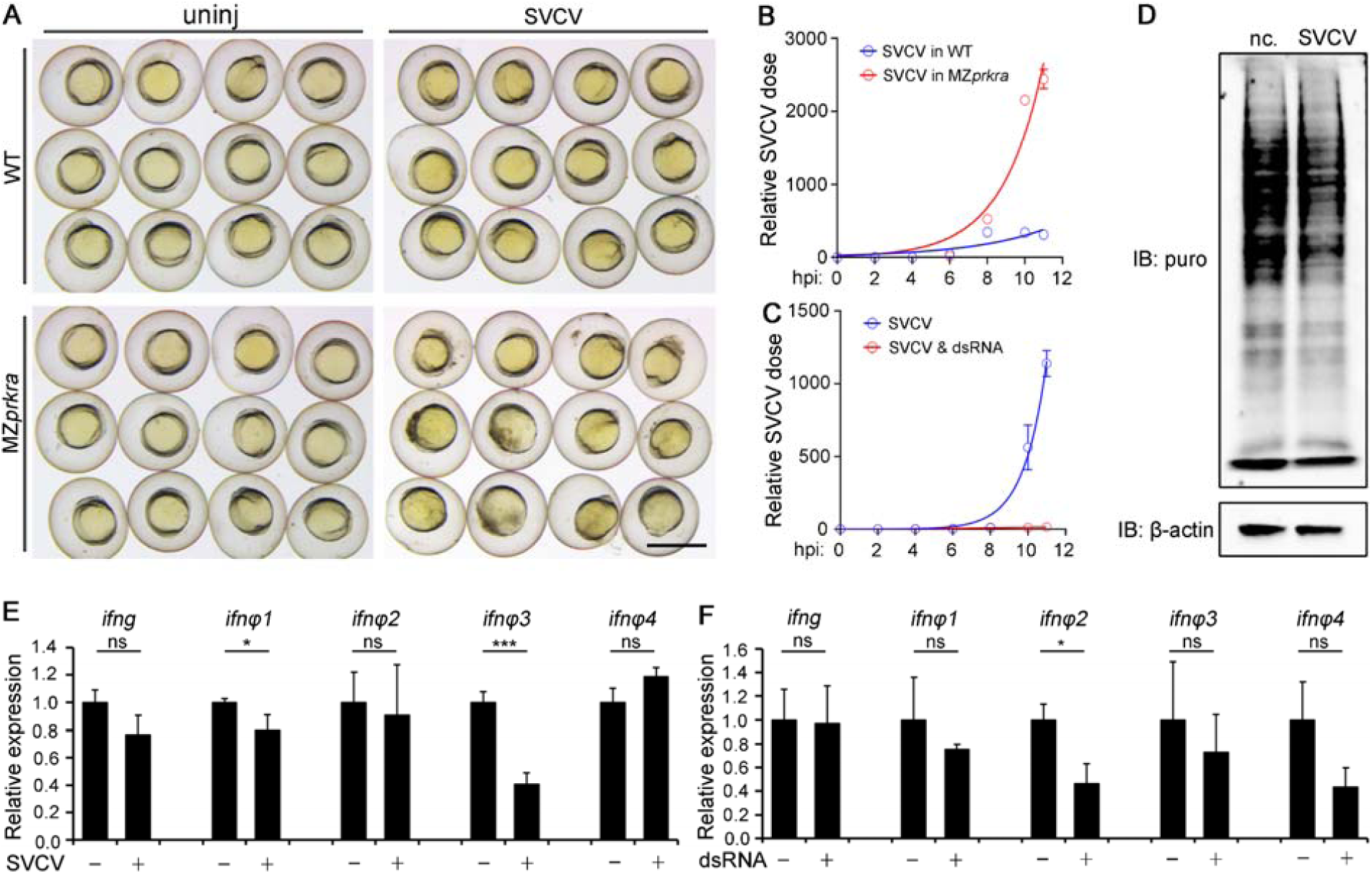
Anti-viral activity of Prkra in zebrafish embryos. (**A**) SVCV infection results in pronounced developmental delays and malformations in MZ*prkra* embryos at 12 hpf. (**B**) Enhanced replication of SVCV in MZ*prkra* mutant embryos. hpi: hours post-injection. (**C**) Suppression of SVCV replication by coinjecting dsRNAs (50 pg/embryo) into wild-type embryos. (**D**) Global reduction of protein synthesis induced by SVCV in the early embryo. (**E**) Failure of SVCV to activate the expression of zebrafish IFN ligand genes in the early embryo. (**F**) Injection of dsRNAs (50 pg/embryo) is also unable to stimulate the expression of IFN genes in zebrafish embryos. Values shown are means ± SDs. * p<0.05, *** p<0.001, ns p>0.05, Student’s *t*-test. The data in **B**-**C** are least-squires fitted according to the logistic growth curve. Scale bars: 1 mm.

### Conserved translational control by Prkra in mice

Finally, we sought to determine whether the dsRNA-induced global translation repression is a species-specific phenomenon or a conserved event. To examine this issue, we injected *gfp* reporter mRNA into early mouse embryos, with or without the addition of dsRNAs. As expected, the injection of dsRNAs significantly suppressed overall GFP expression (Figure 6A). Additionally, we observed a 40%-60% reduction in newly synthesized proteins around E4.5 in response to dsRNA stimulation, as measured by the puromycin incorporation assay (Figures 6B-6D). Consistent with previous reports indicating that dsRNAs do not induce the expression of IFN ligands,^14^ the injection of 100 ng dsRNAs into each embryo failed to upregulate the expression of tested IFN ligand genes, including *Ifna4*, *Ifnab*, *Ifne*, *Ifng*, *Ifnk* (Figure S7A).

**Figure 6.**
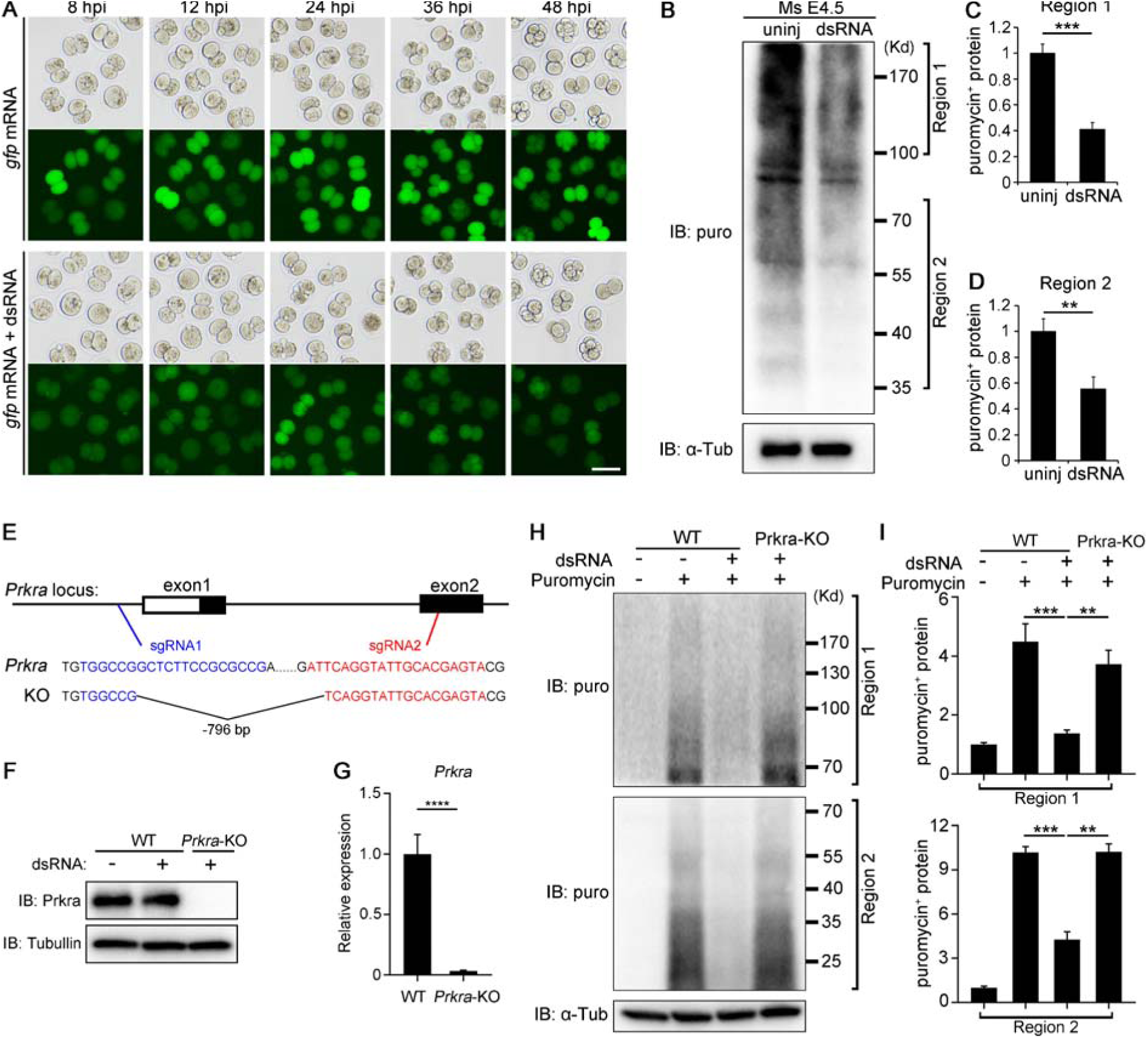
Conservation of Prkra-mediated translation repression in mouse early embryos and mESCs. (**A**) dsRNAs (100 pg/embryo) significantly reduce the translation of *gfp* mRNAs in early mouse embryos. hpi: hours post-injection. (**B**) Inhibition of puromycin incorporation in newly synthesized proteins by dsRNA stimulation (100 pg/embryo). (**C** and **D**) Quantification of puromycin-positive proteins in regions 1 and 2. (**E**) Generation of *Prkra* deficient mESCs. (**F**) Absence of Prkra in *Prkra* deficient mESCs. (**G**) Severely reduced Prkra mRNA in the knockout cell line detected by qRT-PCR. (**H**) Robust translation inhibition by dsRNAs in wild-type mESCs but not in *Prkra* mutants as revealed by puromycin incorporation assay. (**I**) Quantification of puromycin-positive proteins in regions 1 and 2 of **H**. Values shown are means ± SDs. * p<0.05, ** p<0.01, *** p<0.001, **** p<0.0001, ns p>0.05, Student’s *t*-test. Scale bar: 100 μm.

The conservation of dsRNA-induced translational reprogramming was further confirmed in mESCs. To analyze the role of mouse Prkra in translational control, we generated a Prkra knockout (Prkra-KO) mESC line with a large deletion encompassing the entire exon1, intron1 and part of exon2 (Figure 6E). Western blot and qRT-PCR analyses confirmed the absence of Prkra protein and a trace level of transcripts in the mutant cell line (Figures 6F and 6G). Importantly, the knockout of *Prkra* in mESCs nearly abolished the translational inhibitory effect of dsRNAs, as demonstrated by puromycin incorporation assays (Figures 5H and 5I), suggesting a central role for Prkra in mammalian pluripotent stem cells. Nevertheless, dsRNA stimulation failed to induce the expression of *Ifnb1*, *Ifne* and *Ifnk* in mESCs (Figure S7B), excluding the role of IFN system and its downstream PKR in the translational control. These results indicate that Prkra is a conserved key dsRNA sensor and translation regulator in the early embryos and stem cells of vertebrates.

## Discussion

This study uncovered a previously unrecognized mechanism controlling global translation efficiency in vertebrate early embryos and stem cells. These undifferentiated or pluripotent cells utilize Prkra dimer as an independent dsRNA sensor, and dimerized Prkra is activated by dsRNAs to sequester eIF2 complexes from the translation preinitiation complex. This is completely different from the PKR-dominated translational control observed in differentiated cells.

Our findings provide two lines of evidence demonstrating that PKR expression is too low to function effectively in pluripotent cells. First, a low dose of dsRNA injection, which is sufficient to induce significant translation inhibition, does not lead to eIF2α phosphorylation. Second, although high doses of dsRNAs can cause eIF2α phosphorylation, the effect is merely a phenomenon without functional consequence. This is because, in PKR mutants where eIF2α phosphorylation is absent, translation inhibition still persists in response to dsRNAs. Therefore, although Prkra was first identified as a protein activator of PKR,^17,18,20^ its function appears to be independent of PKR in the early embryo.

Prkra binds to the C-terminal domain of RIG-I to facilitate IFN expression in differentiated cells.^23^ However, in stem cells, Prkra is irrelevant to RIG-I-mediated IFN signaling. Our data show that neither dsRNA nor SVCV induces the upregulation of IFN ligand gene expression in early zebrafish embryos. In mouse embryos and mESCs, dsRNA stimulation also fails to upregulate IFN ligand gene expression. These observations are consistent with previous findings that ESCs and embryonic carcinoma cells do not produce type I IFNs in response to viral infection or poly(I:C) treatment.^14^ Hence, these data rule out a functional connection between Prkra and RIG-I in early embryos.

It has been reported that the dsRBD3 of Prkra binds to the Partner-binding domain (PBD) of Dicer,^32^ making the role of Prkra controversial in strand selection during RNAi and siRNA maturation.^32,39^ However, in early embryos, the function of Prkra in microRNA production may not be essential for normal development. Although we did not test the status of miR430, the only essential microRNA in early development of zebrafish^40^, in MZ*prkra* mutant embryos, the normal development of *prkra* mutants strongly suggests that miR430 may still be functional.

Therefore, our findings provide critical evidence that Prkra-mediated translation repression operates independently of PKR in undifferentiated cells, and that its function is also independent of Rig-I and Dicer. In fact, the regulation of PKR and Dicer activity is mediated by Prkra monomers.^30,32^ However, while previous studies have demonstrated that Prkra forms dimers,^31,33^ the function of these dimers remains unknown. We propose that Prkra dimers act as critical dsRNA sensors and present a model elucidating the molecular mechanism by which Prkra activation mediates global translation inhibition. Our biochemical analysis indicates that Prkra dimers bind to dsRNAs in a highly organized manner, as evidenced by the ordered bands in EMSA analysis. The dimerized dsRBD3 is essential for Prkra to sequester the eIF2 complex. According to Alphafold3 modeling, the dsRBD3 dimers may be neatly arranged on one side of the dsRNA, a structure suggesting that two or more adjacent dsRBD3 dimers is likely to mediate the binding to the eIF2 complex (Figures 7A and 7B). Thus, dsRNA-activated Prkra grasps the eIF2 away from the preinitiation complex essential for translation initiation, leading to significant reduction of protein synthesis (Figure 7B). Structural biology analysis will be beneficial to further verify this promising model.

**Figure 7.**
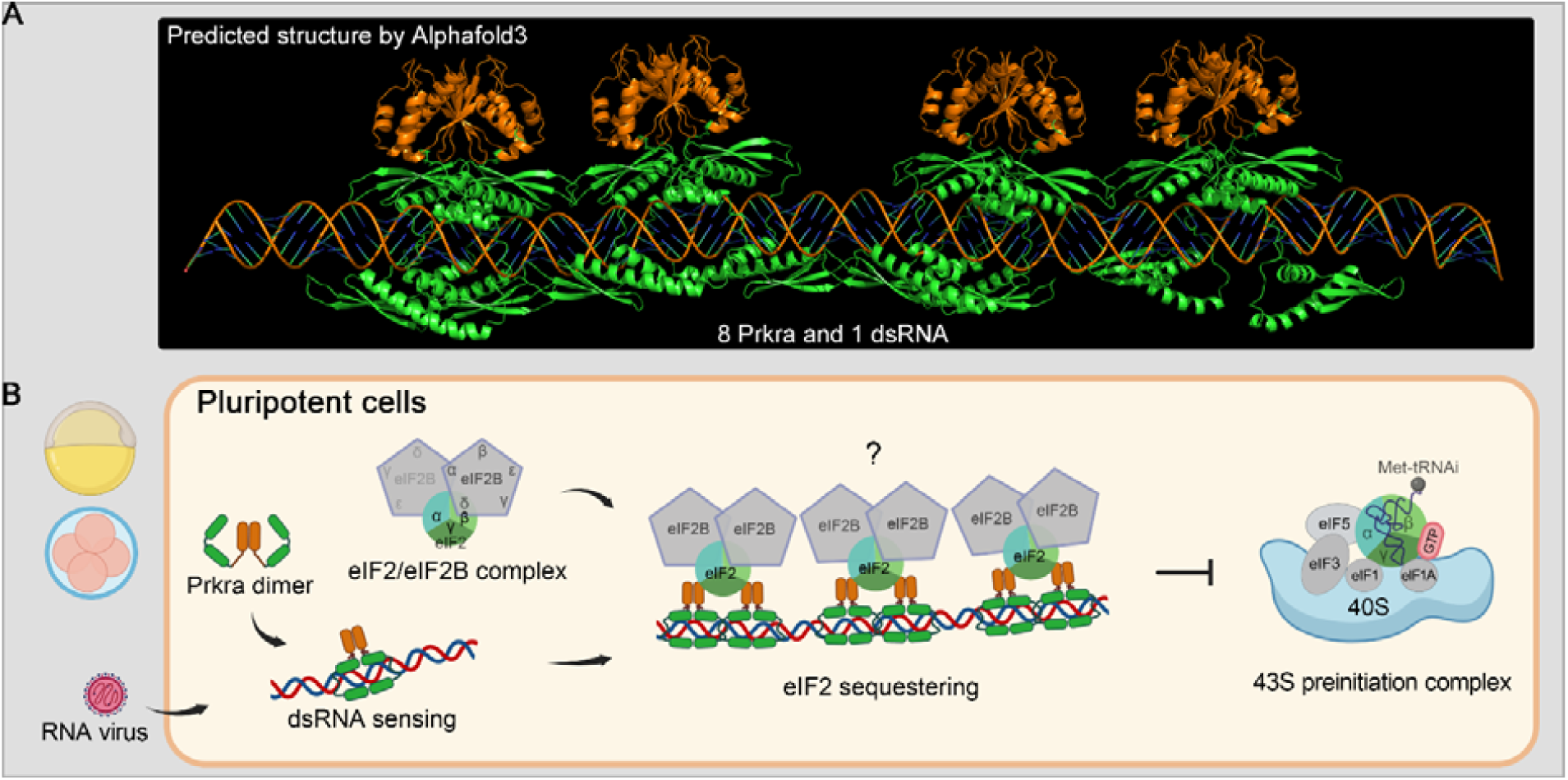
Model of Prkra dimer-mediated dsRNA sensing and translation control in pluripotent cells. (**A**) Predicted structure of eight Prkra proteins bound to a 139 bp dsRNA, as modeled by AlphaFold3. The dimerized dsRBD3s are highlighted by orange. (**B**) Prkra exists as a dimer that senses and aligns with dsRNA in a highly ordered manner. Utilizing its dimerized dsRBD3s, the Prkra-dsRNA complexes interact with and sequester eIF2, potentially in conjunction with eIF2B, thereby hindering the formation of the active eIF2 complex necessary for the assembly of the 43S preinitiation complex, which is critical for efficient translation initiation.

The robust stress response induced by dsRNAs highlights the challenges of employing siRNA-mediated knockdown in animal models such as zebrafish and *Xenopus*.^41,42^ Since the discovery of RNA interference in the late 1990s, dsRNAs have been introduced into zebrafish eggs.^43^ However, dsRNA injection induced toxic effects rather than specific gene knockdown, limiting the application of siRNA in zebrafish.^34,42^ It was hypothesized that dsRNAs could interfere with the biogenesis and activity of the essential endogenous miRNA, miR430, and induce an IFN-mediated innate immune response in zebrafish embryos.^44,45^ Our study excludes these possibilities and ultimately resolves the longstanding enigma. We demonstrate that dsRNAs induce strong global translation inhibition, resulting in toxicity to early zebrafish embryos. In contrast, mouse embryos may exhibit lower sensitivity to dsRNA-induced translation repression or possess higher RNA interference efficiency than zebrafish, making siRNA-mediated knockdown feasible in this context. Nevertheless, avoiding excessive siRNA injection into early mouse embryos may be crucial to prevent nonspecific toxic effects associated with Prkra-mediated translation repression.

In conclusion, our study identifies Prkra dimer as a dsRNA sensor in embryonic and stem cells, which plays a crucial role in regulating an anti-viral global translation inhibition. Future research should aim to elucidate the extent to which Prkra-mediated translational control compensates for the PKR-eIF2α axis in differentiated cells. Moreover, investigating the potential of this translation inhibition mechanism in the development of novel antiviral and anti-tumor therapies would be of significant interest.

## Supporting information

Data S3 RNA-pulldown followed by MS, metafile

Data S1 Ribo seq, metafile

Data S2 Gene expression related to this manuscript

Uncropped Blots

Raw data for each Figure

## Data availability

The deep sequencing data of Ribo-seq are available from NCBI Gene Expression Omnibus GSE234230. The scripts are available at https://github.com/benjaminfang/ribo-seq-workflow. Further information and requests for resources and reagents should be directed to and will be fulfilled by the corresponding author, Ming Shao.

## Supplementary Data statement

Figures S1-S6.

Tables S1 and S2.

Data S1. (separate file) Metadata spreadsheet of Ribo-seq data.

Data S2. (separate file) Expression of genes mentioned in this study.

Data S3. (separate file) Metadata spreadsheet of dsRNA pulldown followed by LC-MS/MS.

Data S4. (separate file) Raw data visualized in each Figure.

Data S5. (separate file) Uncropped blots presented in this paper.

## Acknowledgements

We are grateful to professors Anming Meng and Li Yu for sharing their ideas for improving the work. We thank J.L.S and H.J.D from the Core Facility and Service Platform, School of Life Sciences, H.Y.Y and X.M.Z from SKLMT for assistance with confocal imaging, histological sectioning and ribosome profiling, and Z.X.D for fish rearing.

## Supplementary Figures

**Figure S1.**
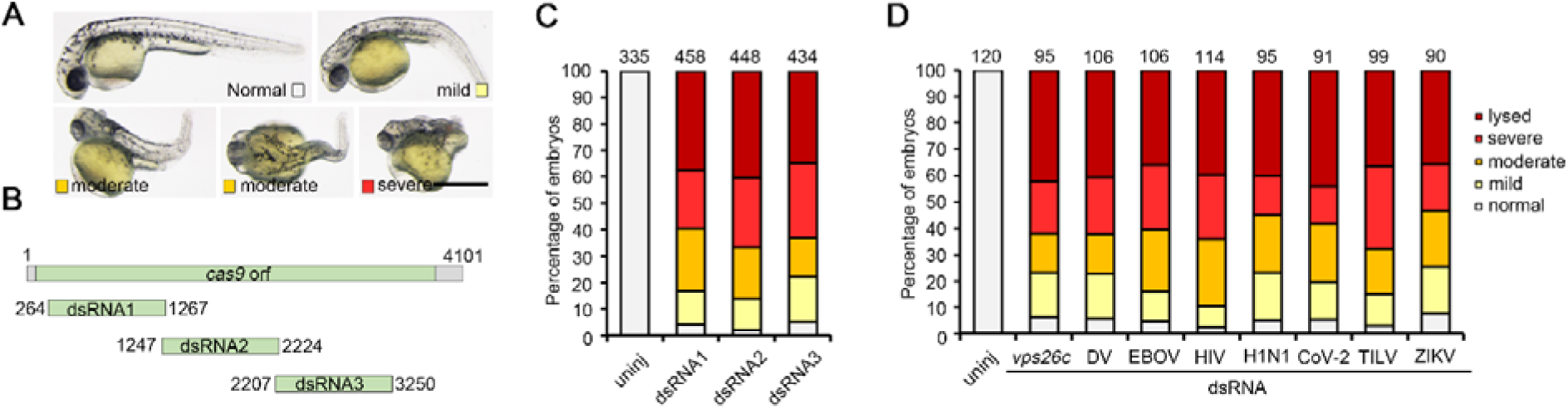
dsRNAs induce toxic effects on early development in a sequence-independent manner. (**A**) Representative toxic phenotypes caused by dsRNA injection. (**B**) 1-kb dsRNAs derived from sequences of *cas9* orf. (**C** and **D**) Phenotypes caused by injection of dsRNAs from different sequence sources (see Table S1). Scale bars: 0.5 mm.

**Figure S2.**
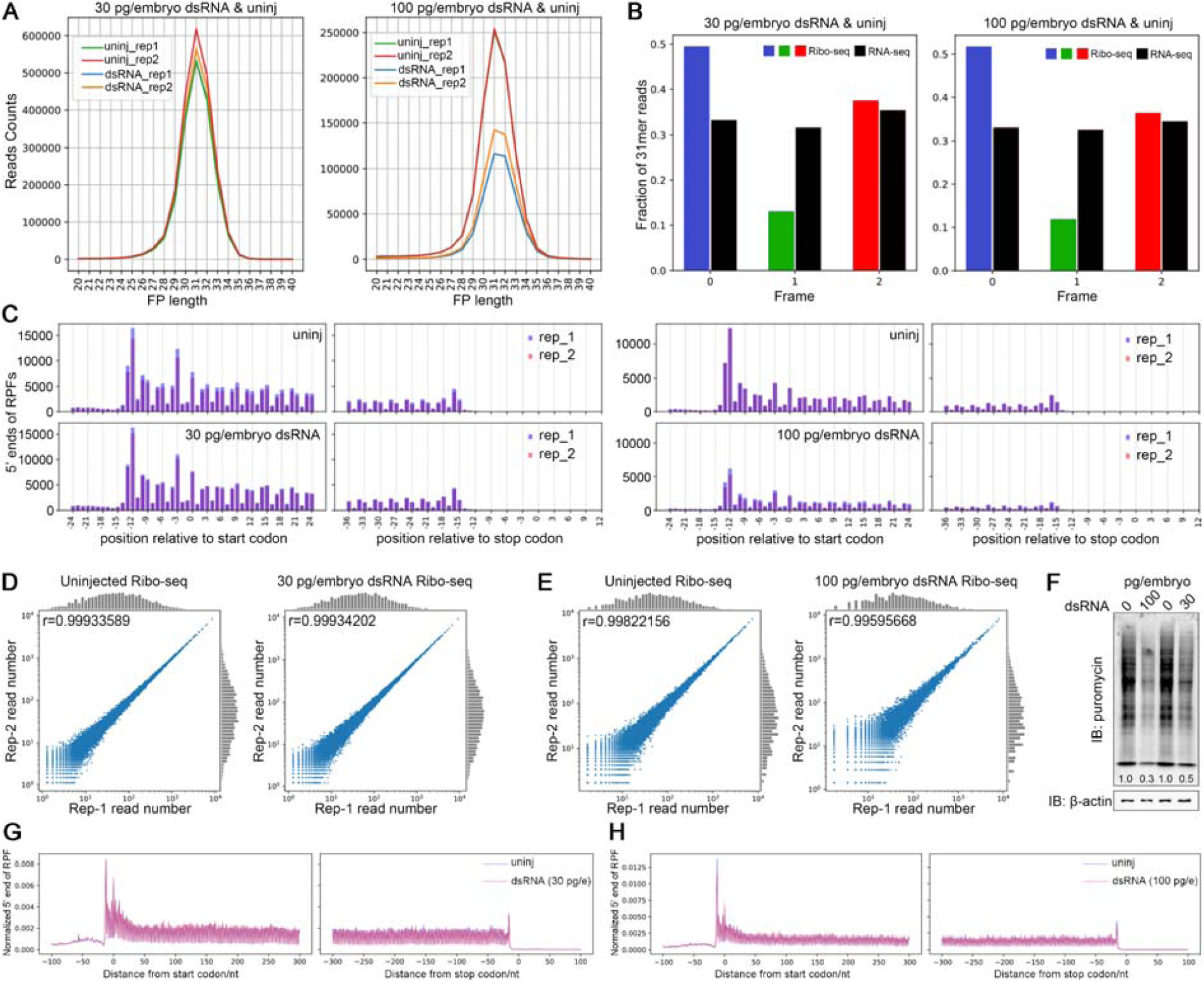
Ribo-seq analysis of translation remodeling after dsRNA stimulation. (**A**) Read-length distribution of uniquely mapped RPFs. (**B**) Triplet periodicity of ribosome footprints revealed by reads aligning to the A-site of each reading frame but not by reads from RNA-seq. (**C**) Metagene analysis of ribosome profiling libraries, with 5’ end of reads aligned to the start or stop codon. (**D** and **E**) Scatter plots illustrating the reproducibility between replicates of Ribo-seq. (**F**) A representative puromycin incorporation experiment demonstrating the degree of reduction in global translation activity. (**G** and **H**) 5’ end distribution of ribosome footprints on mRNAs from uninjected and dsRNA-injected embryos.

**Figure S3.**
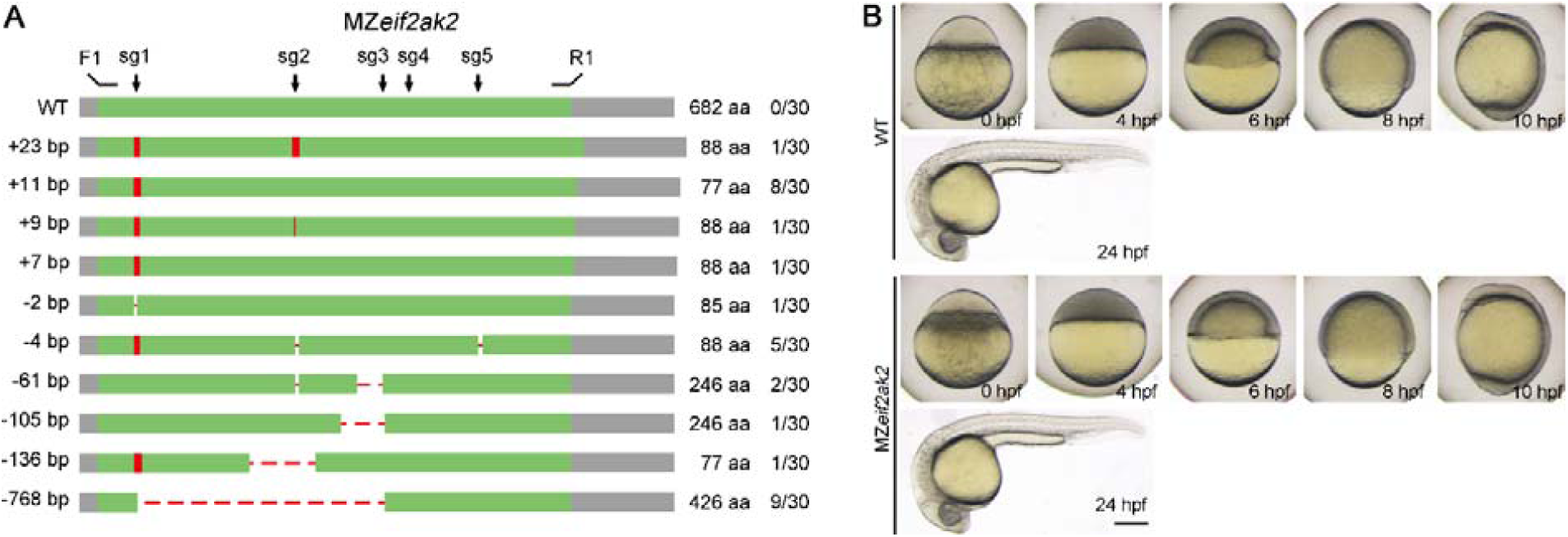
Genotyping and phenotyping of PKR maternal mutants. (**A**) Injection of Cas9 RNPs targeting the ORF of *eif2ak2* into 1-cell stage zebrafish embryos. The CRISPants were raised to adults and crossed to produce MZ mutant embryos. ORF region of *eif2ak2* (green area) was amplified and sequenced from cDNAs of CRISPant-spawned embryos at 6 hpf. The result indicates the absence of wild-type alleles. sgRNA target sites are indicated by arrows. F1 and R1 are primers amplifying the ORF. UTRs are marked as grey. (**B**) Maternal and zygotic mutants of *eif2ak2* develop normally from fertilized egg to 24 hpf. Scale bar: 200 μm.

**Figure S4.**
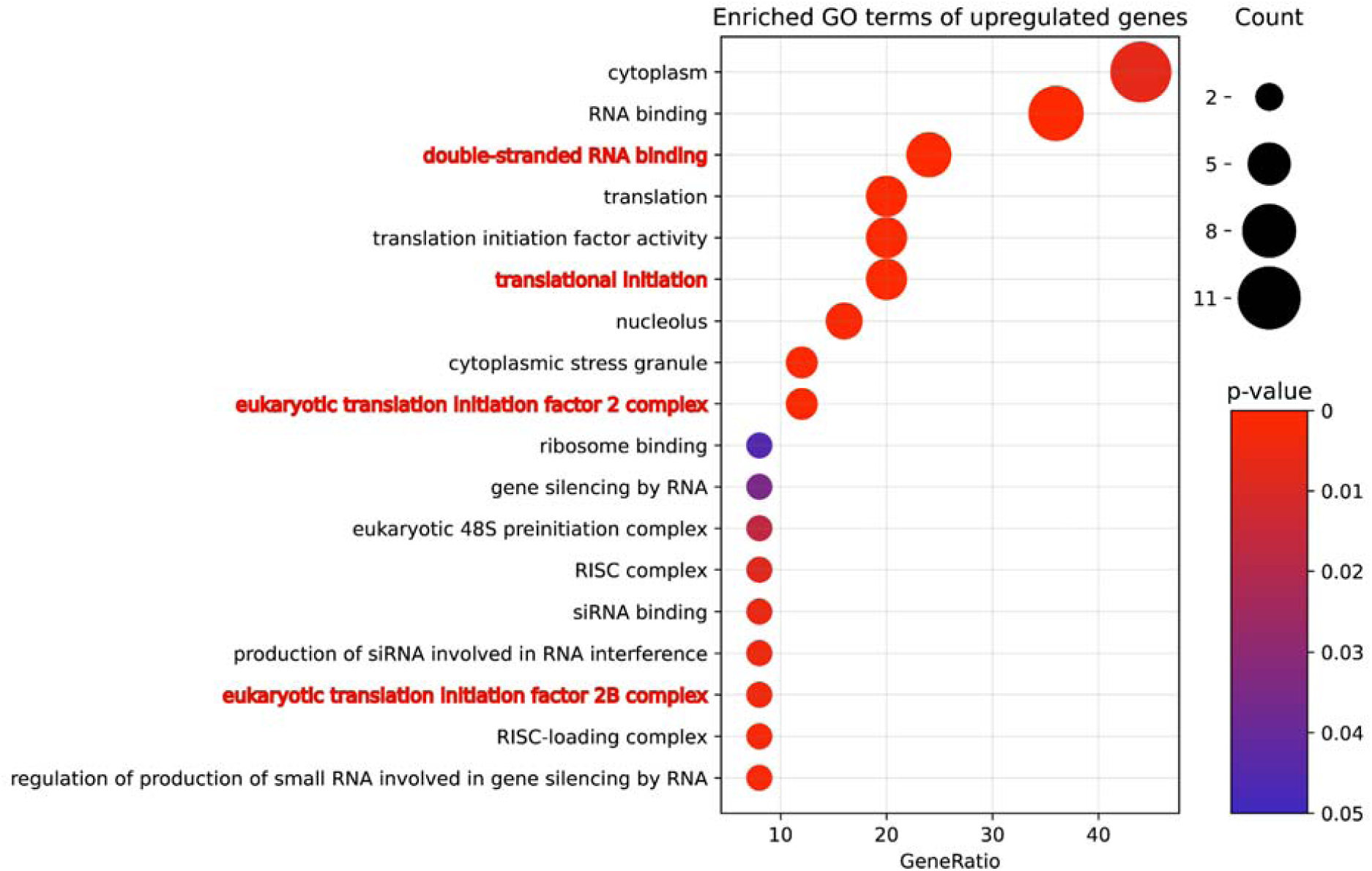
Enriched GO terms of proteins specifically pulled down by dsRNAs. Red highlights dsRNA binding and translation-associated terms.

**Figure S5.**
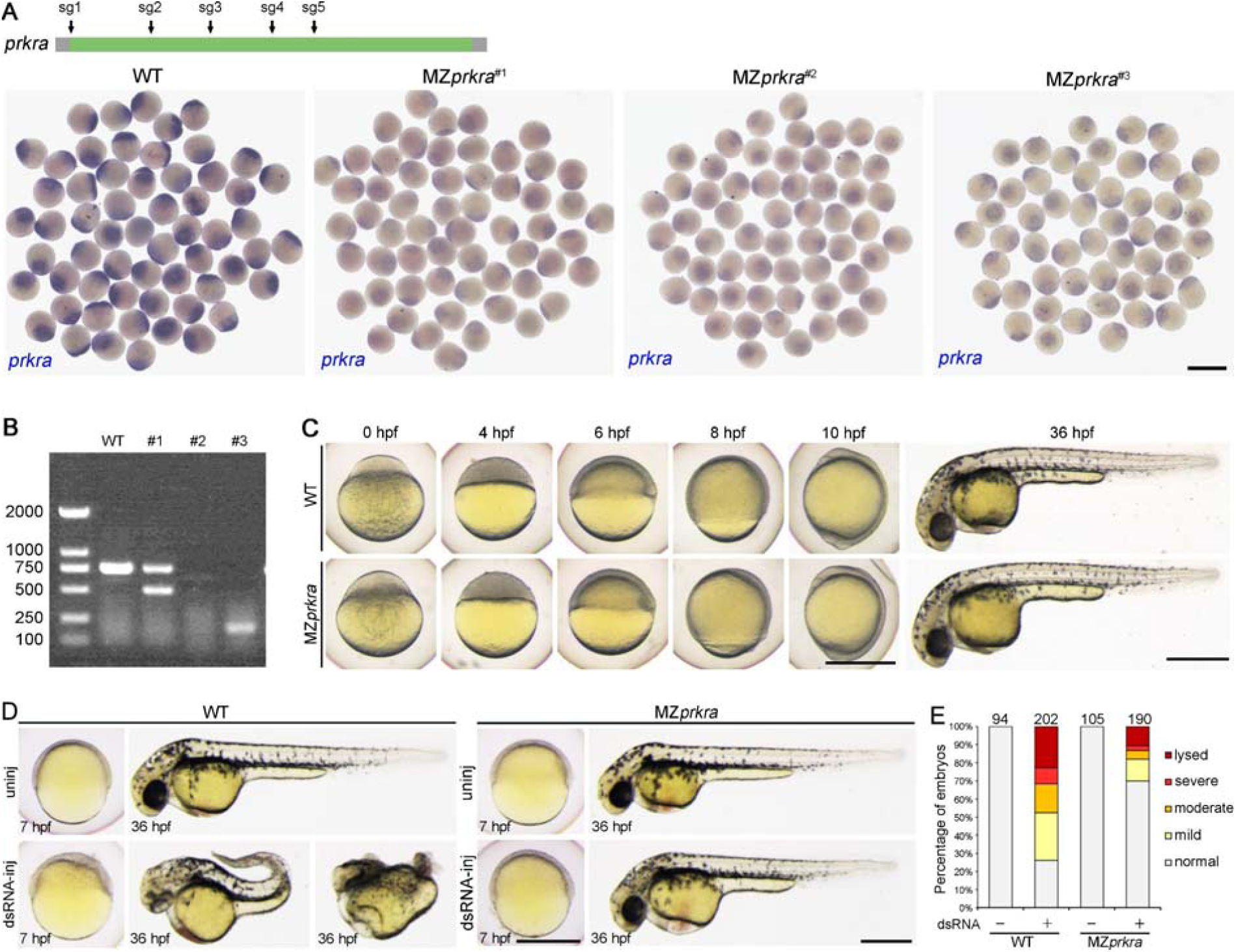
Genotyping and phenotyping of MZ*prkra*. (**A**) MZ*prkra* mutant embryos at the 1-cell stage from three independent F0 females, showing obvious non-sense mediated decay. (**B**) RT-PCR analysis of *prkra* CDS in MZ*prkra* embryos at the 4-cell stage. (**C**) MZ*prkra* mutants do not exhibit defects in early development. (**D**) Representative phenotypes caused by injection of dsRNAs (50 pg/embryo). (**E**) Histogram showing the reduced toxicity of dsRNAs in MZ*prkra* embryos. Scale bars: 1 mm for **A**, 500 μm for **C** and **D**.

**Figure S6.**
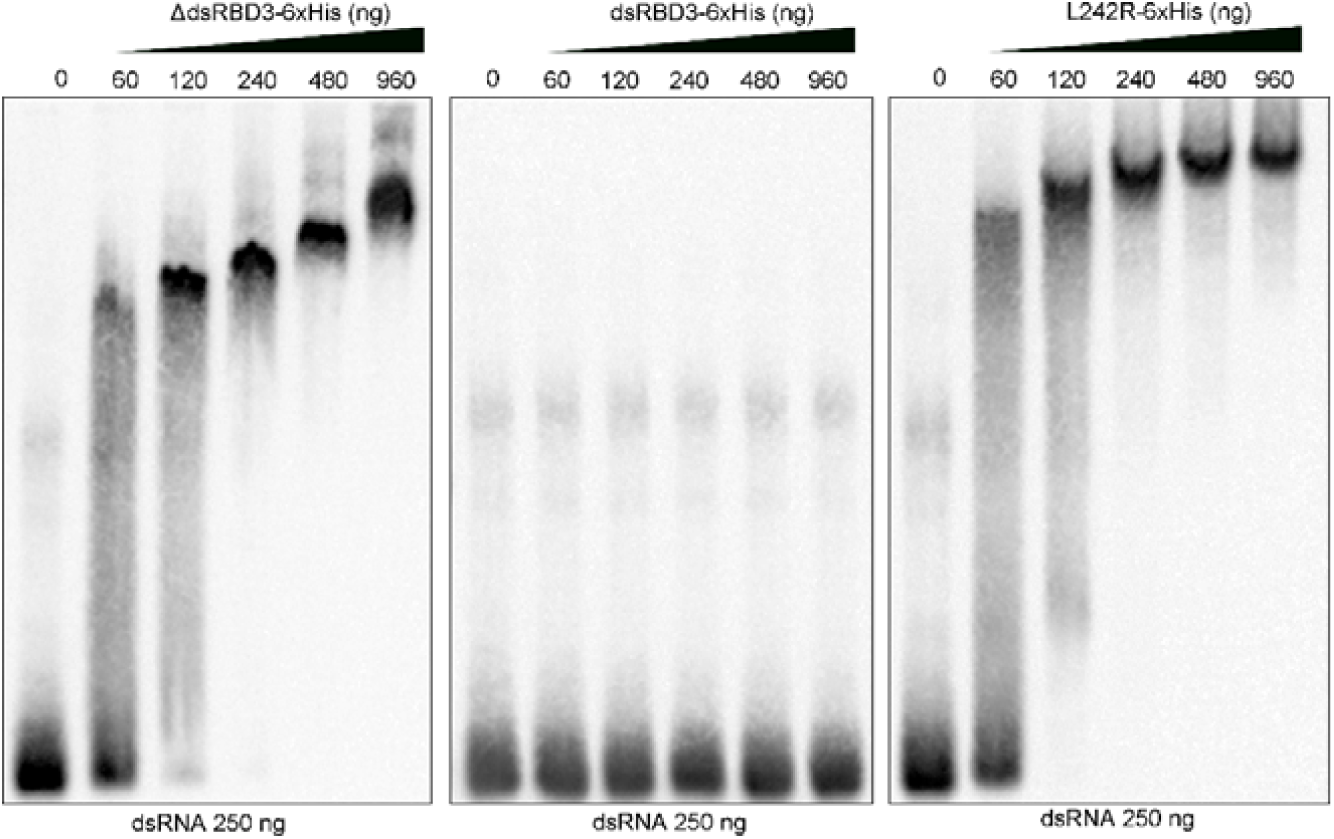
EMSA analysis of dsRNA binding by mutant Prkra proteins. Note that only dsRBD3 fails to bind to dsRNAs, while the dimerization-deficient mutants ΔdsRBD3 and L242R do not produce bands with a laddering pattern in the EMSA gel. Instead, their dsRNA complexes exhibit a smeared appearance when these proteins are added at 60 and 120 ng/lane.

**Figure S7.**
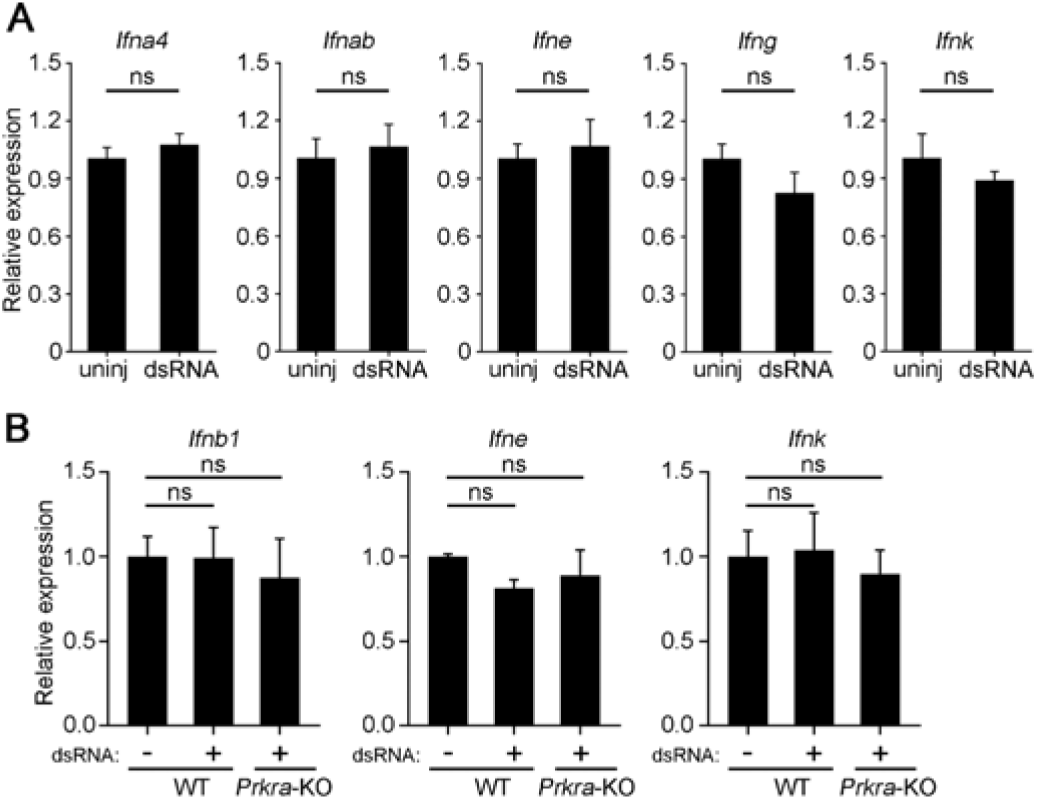
dsRNA stimulation fails to induce the expression of IFN ligand genes in early mouse embryos and mESCs. (**A**) dsRNA injection at 100 pg/embryo has no effect on the expression of IFN ligand genes in early mouse embryos (E4.5). (**B**) IFN ligand gene expression is not induced by dsRNAs in wild-type and *Prkra*-KO mESCs. ns p>0.05, Student’s *t*-test.

## Supplementary Tables

**Table S1.**
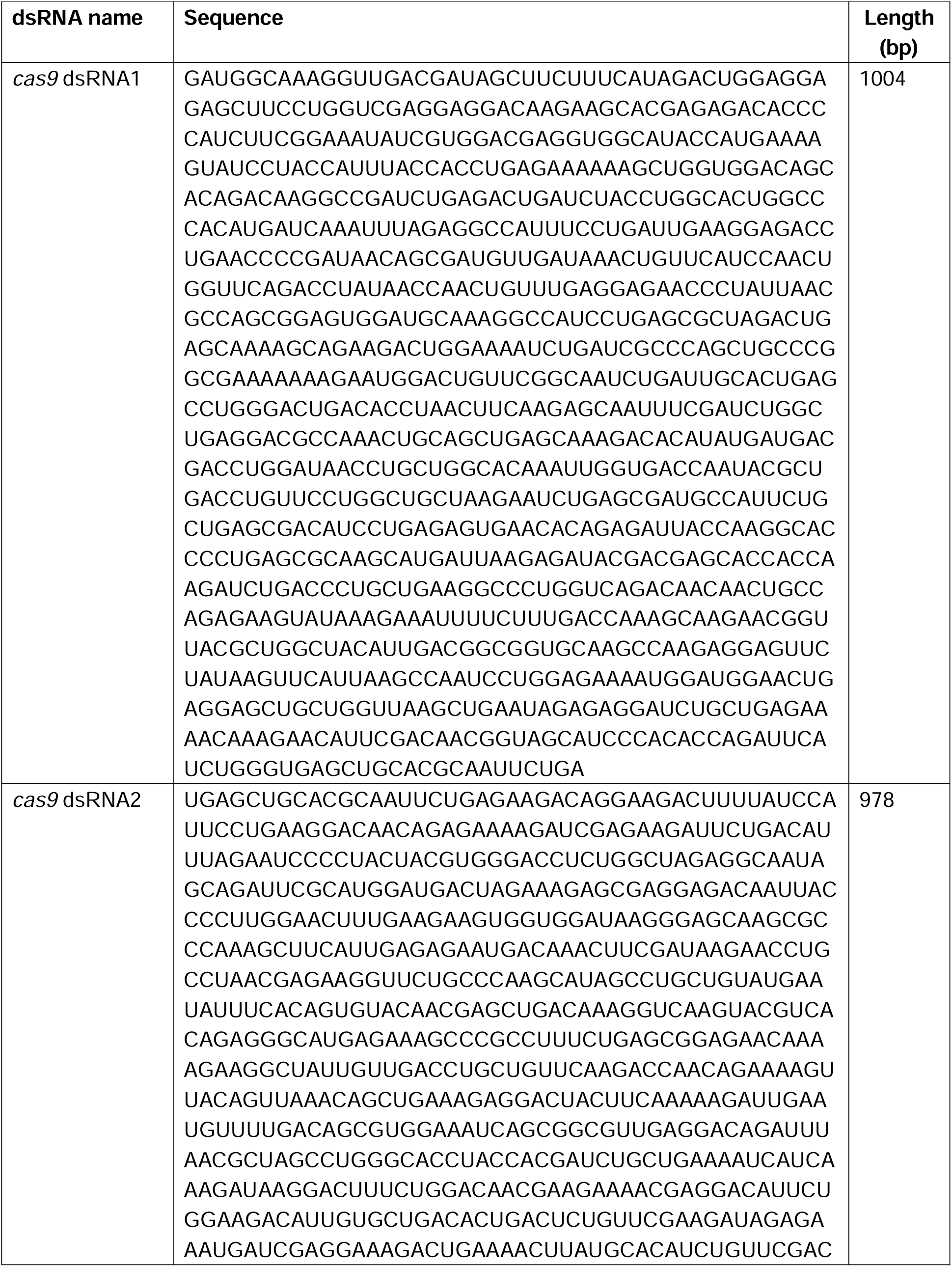
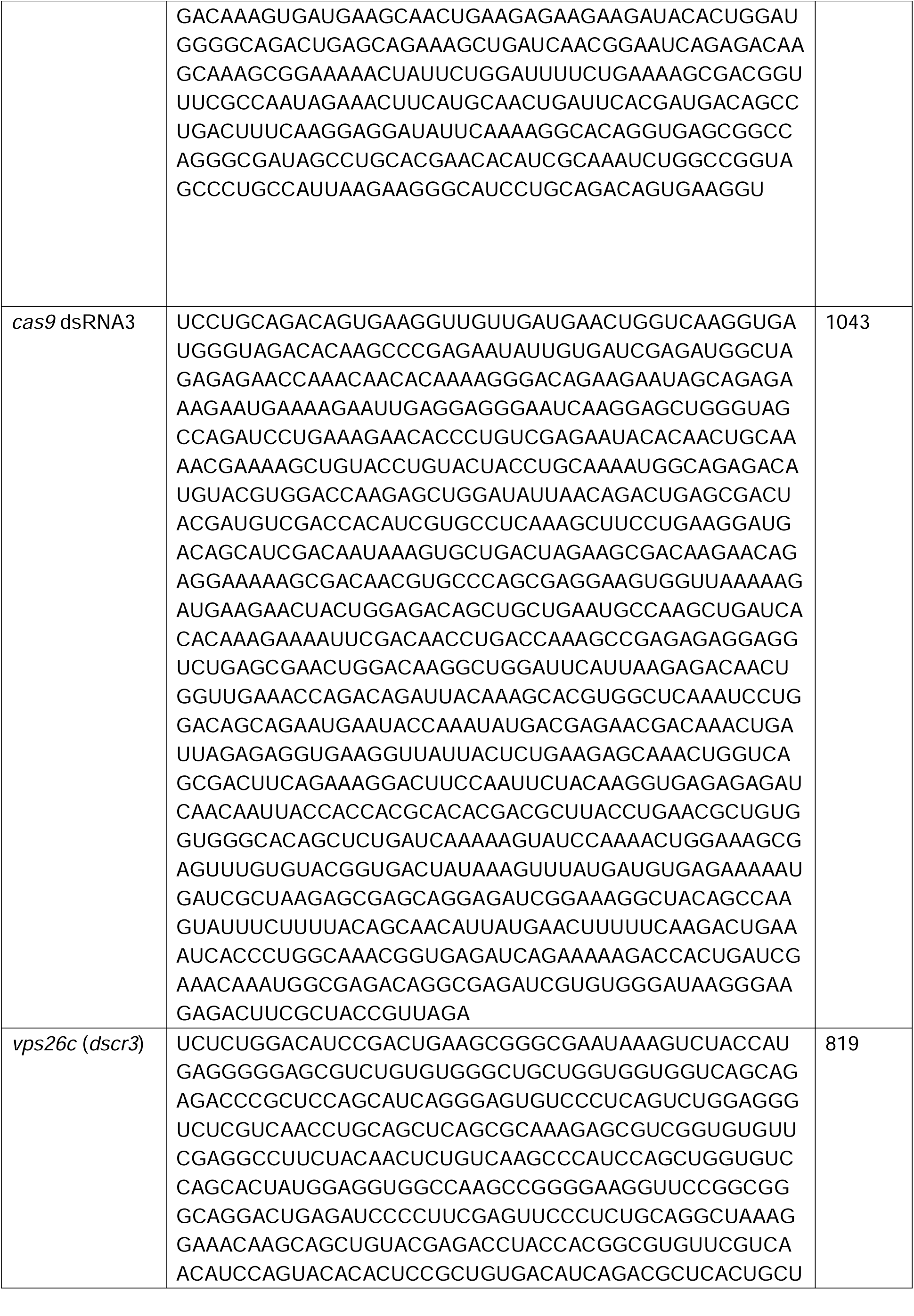
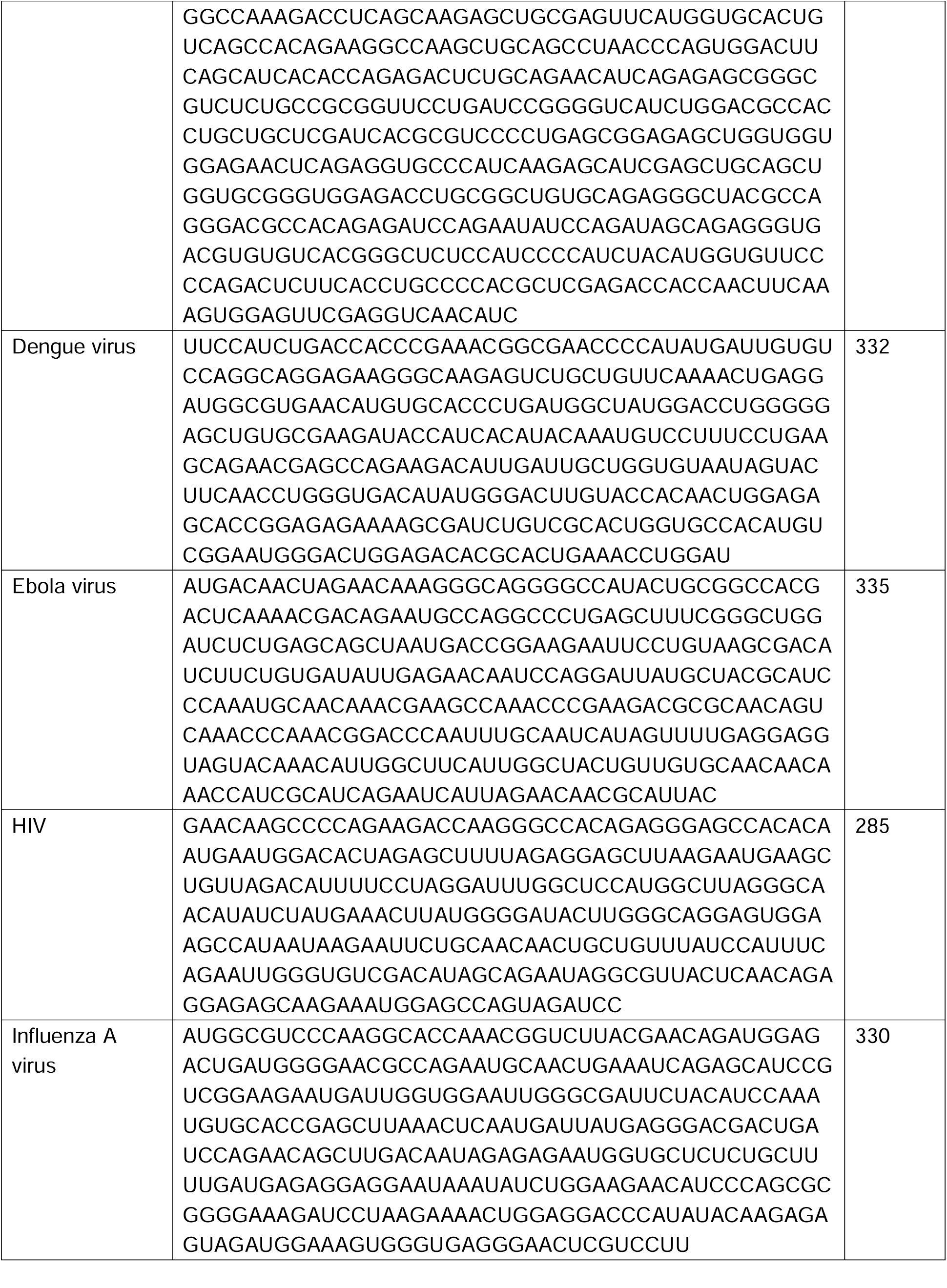
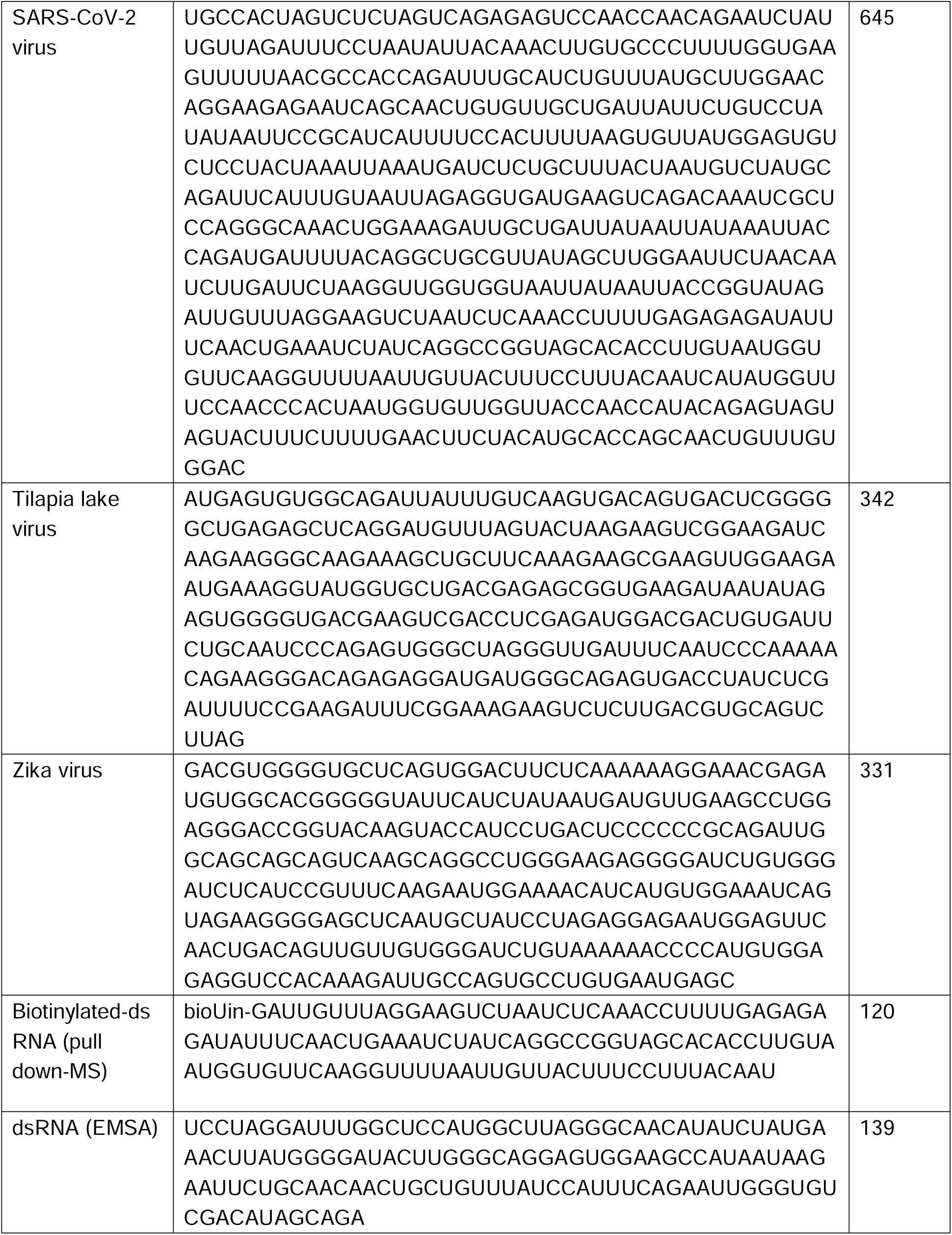
Sequences of dsRNAs.

**Table S2.**
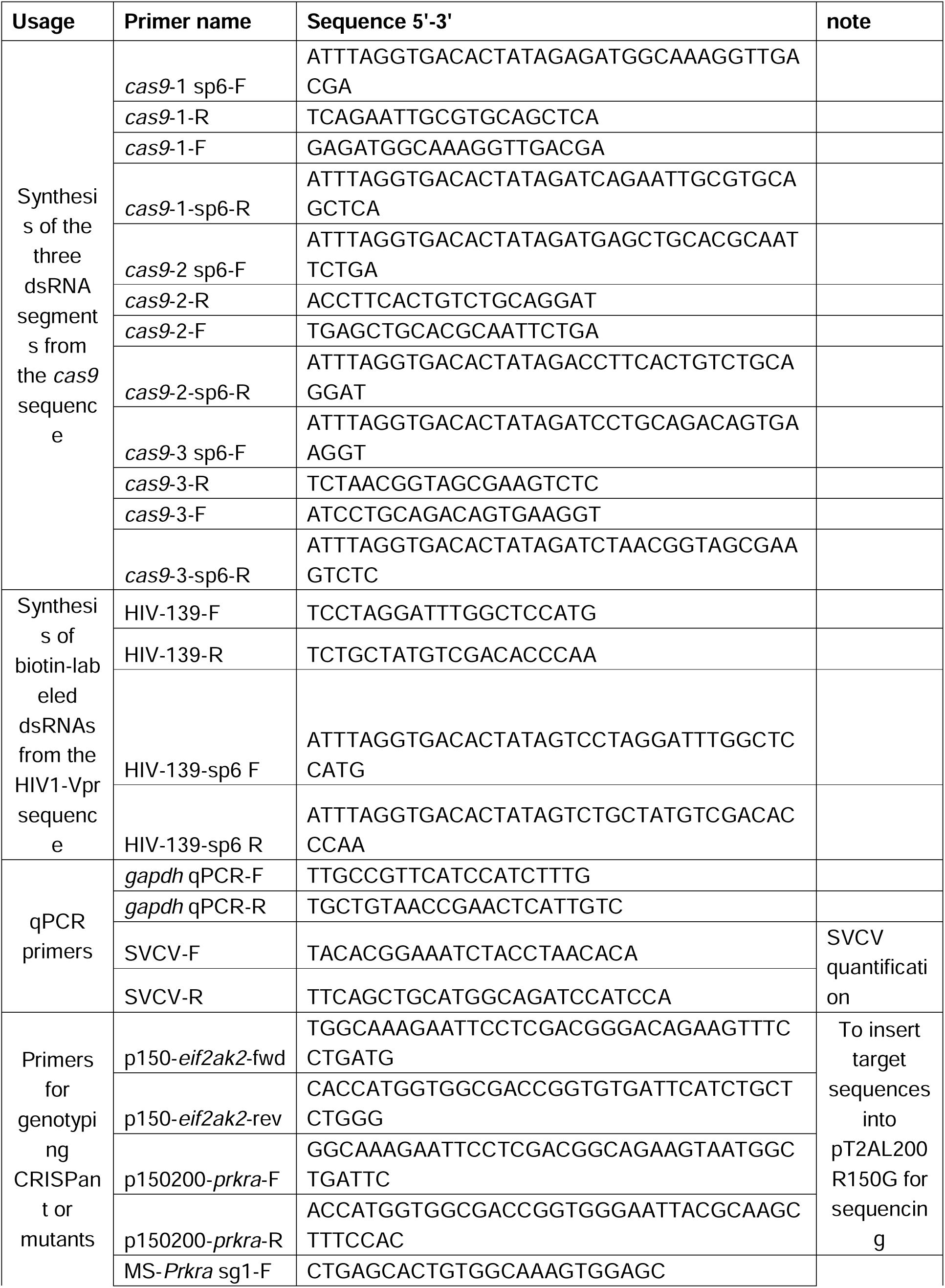
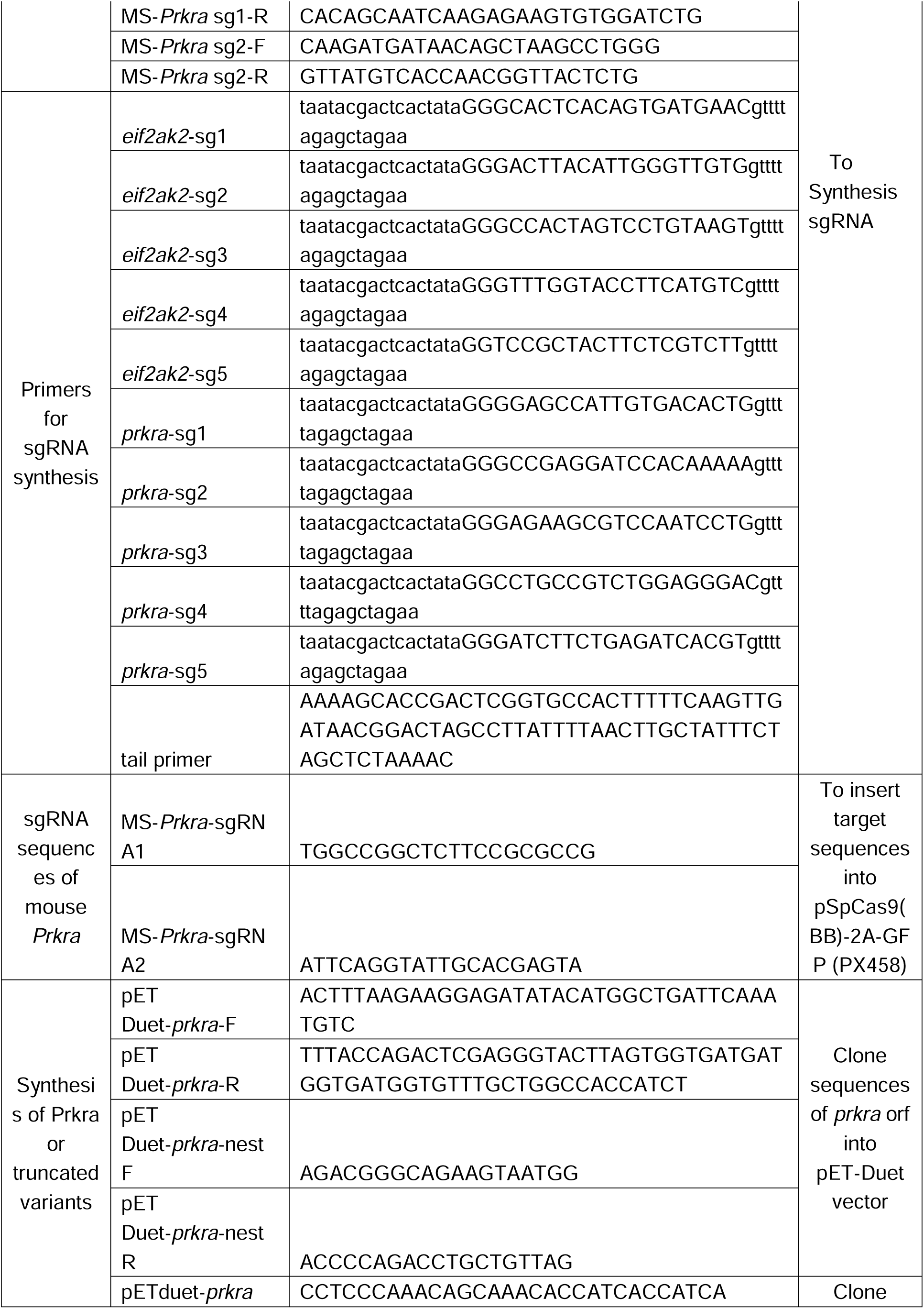
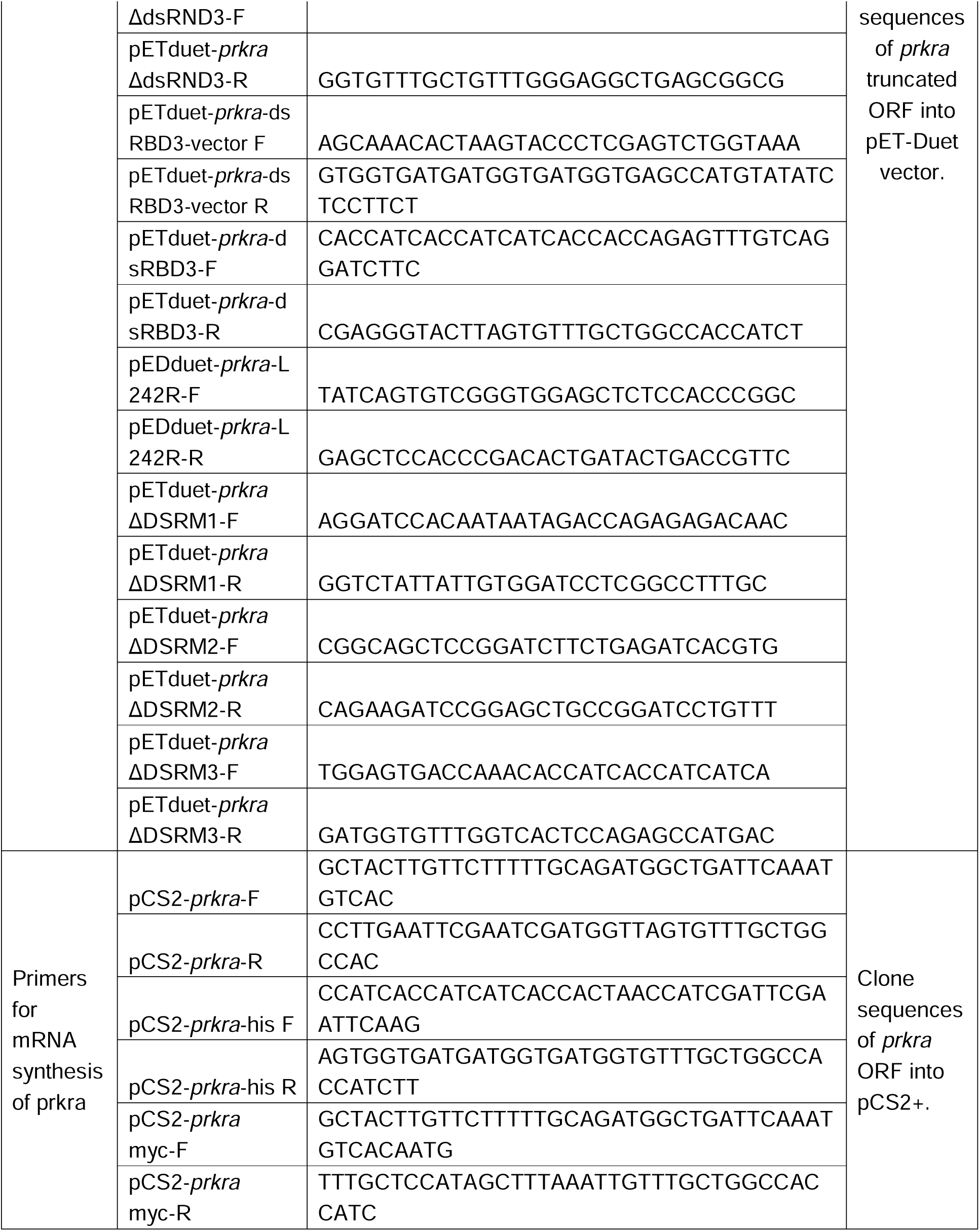
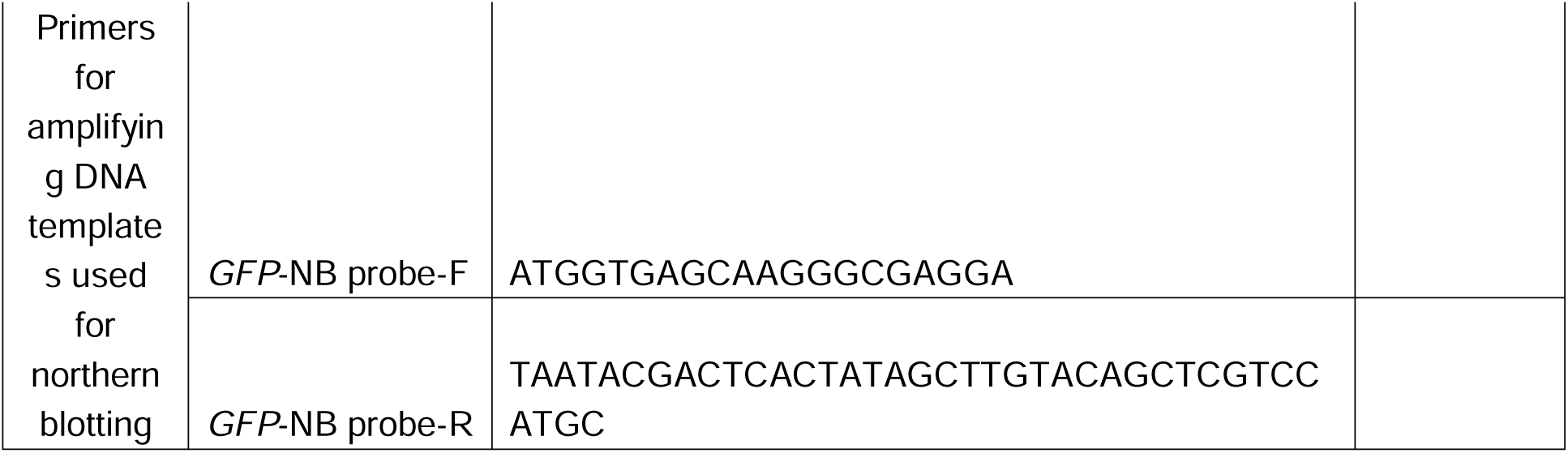
Primers used in this study.

## STAR Methods

### KEY RESOURCE TABLE

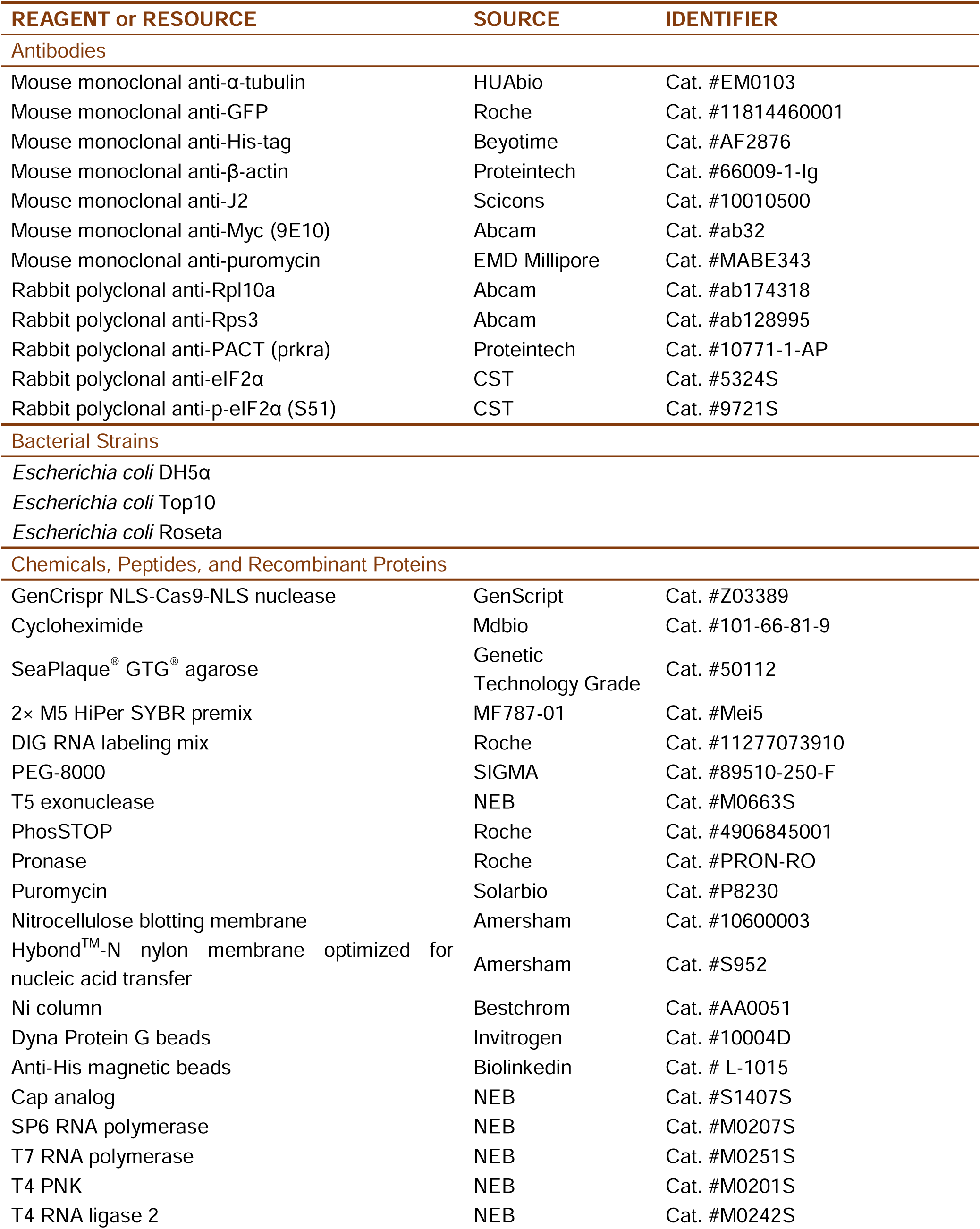
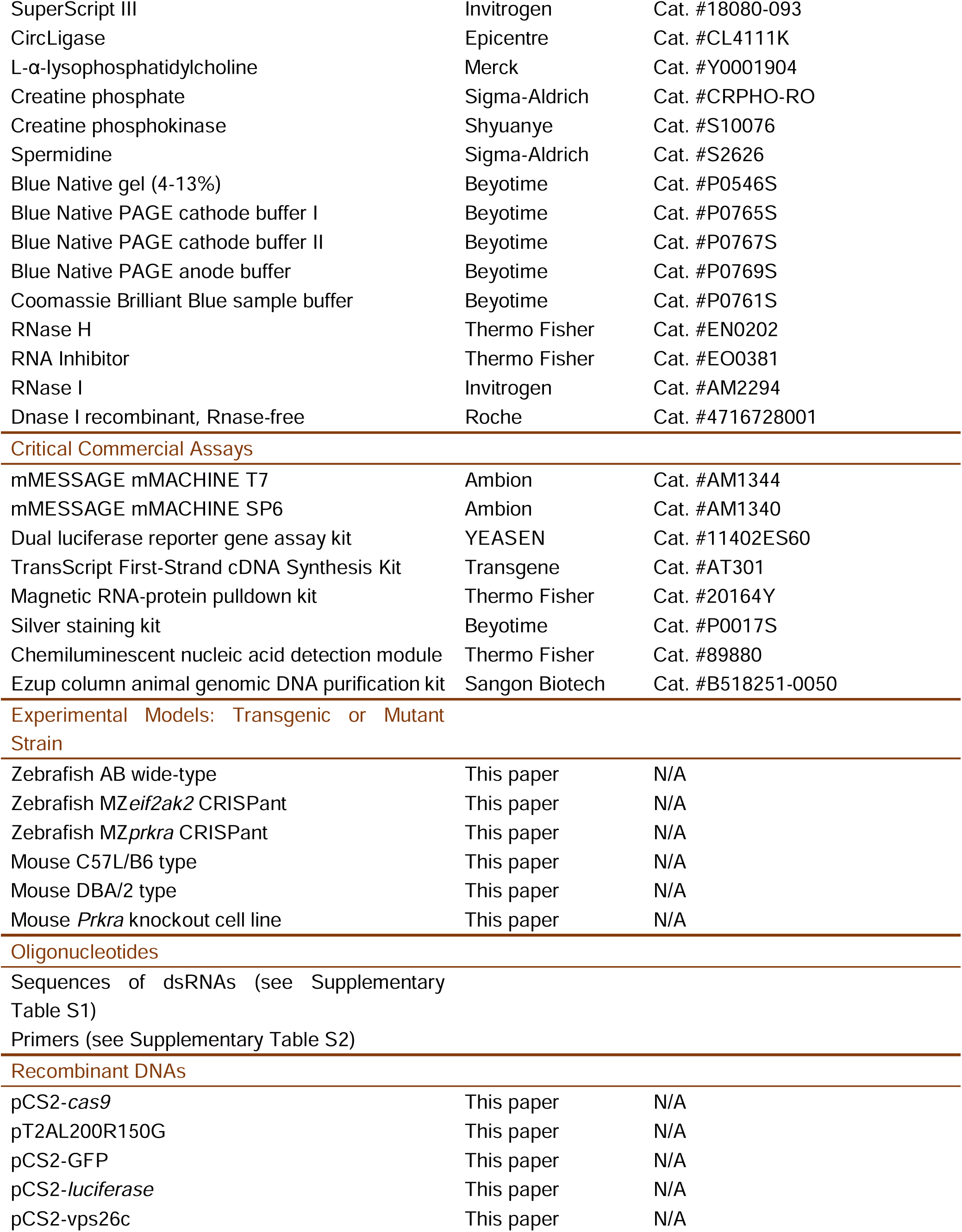
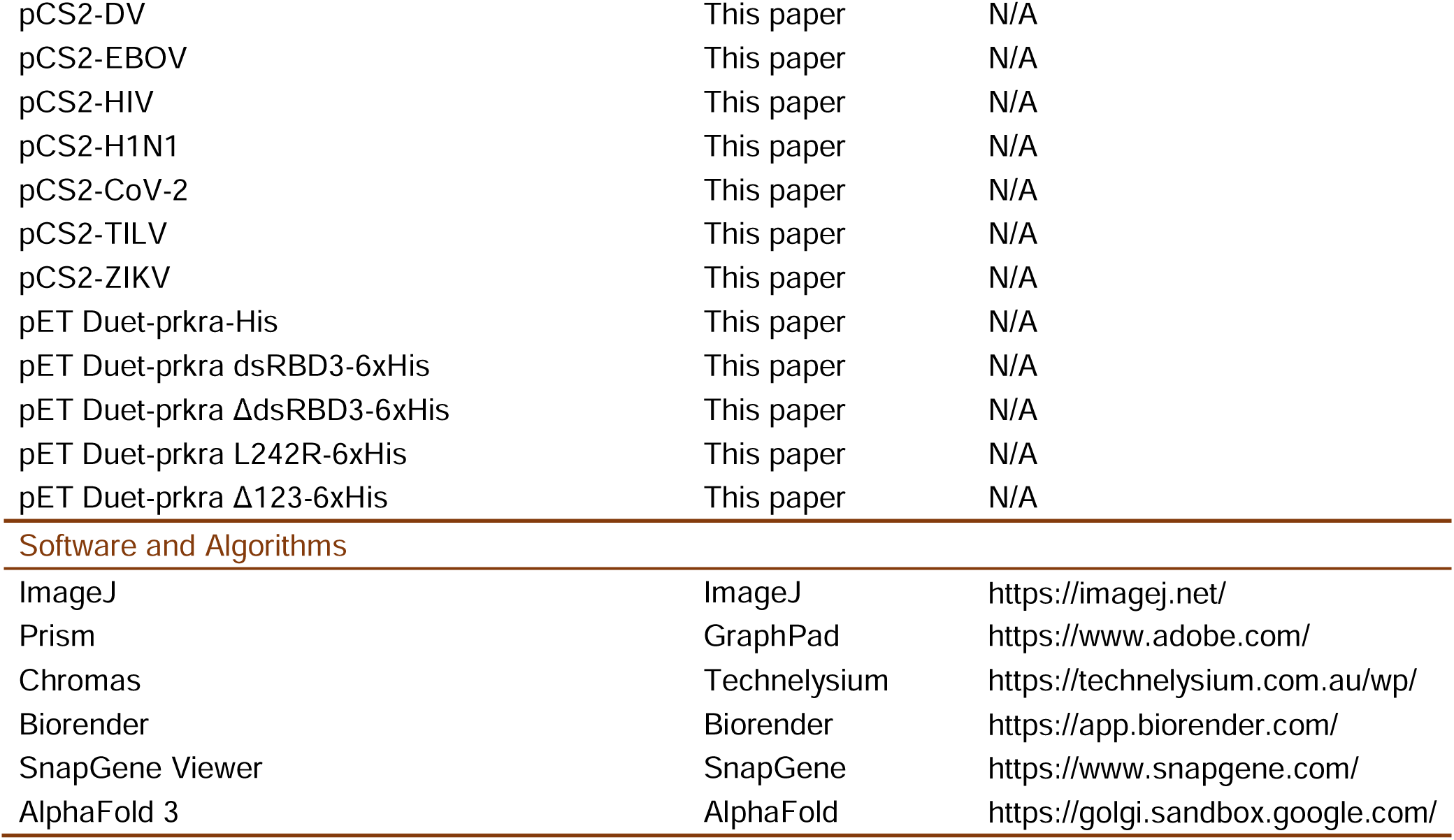

### MATERIALS AVAILABILITY

The deep sequencing data of Ribo-seq and RNA-seq are available from NCBI Gene Expression Omnibus GSE234230 and GSE265771. The scripts are available at https://github.com/benjaminfang/ribo-seq-workflow. Further information and requests for resources and reagents should be directed to and will be fulfilled by the lead contact, Ming Shao.

### EXPERIMENTAL MODELS AND SUBJECT DETAILS

#### Animals

Zebrafish AB strain, *eif2ak2* and *prkra* mutant lines were raised in standard breeding systems (Haisheng) at 28 °C. All experiments followed the ARRIVE guidelines approved by the Ethics Committee for Animal Research of Life Science of Shandong University (permit number SYDWLL-2023-069).

All mice with C57L/B6 and DBA/2 backgrounds were kept in individually ventilated cages (IVC) in a specific pathogen-free (SPF) environment and handled according to guidelines approved by the Animal Care and Use Committee of the Kunming Institute of Zoology, Chinese Academy of Sciences (IACUC-RE-2024-05-002).

### METHOD DETAILS

#### Generation of zebrafish maternal and zygotic mutants

We adopted a CRISPant strategy to generate zygotic mutants and maternal mutants rapidly.^37^ sgRNAs targeting the open reading frames (ORFs) of *eif2ak2* and *prkra* were designed by the online tools CRISPRscan.^46^ They were prepared via standard *in vitro* transcription and purification. The purified sgRNAs (100 pg) were injected with Cas9 protein (200 pg) into the fertilized egg. At 10 hpf, embryos were homogenized in 50 mM NaOH for 20 min at 95 °C followed by the addition of 1/10 volume of 1 M Tris-HCl (pH 7.5). The DNA fragment flanking the sgRNA binding site was PCR amplified and sequenced. Sequencing chromatogram files were submitted to http://shinyapps.datacurators.nl/tide/ to estimate the efficiency of the sgRNA. The adult CRISPant male and female F0 fish were crossed to obtain maternal and zygotic (MZ) mutant embryos, which were verified by RT-PCR analysis using cDNAs from 1-cell stage embryos for analysis of maternal transcripts or from shield-stage embryos for analysis of MZ transcripts. The PCR products were also subcloned and subjected to Sanger sequencing.

#### Mouse embryos and mESC knockout lines

Female C57BL/6 (6-8 weeks old) were induced to superovulation by intraperitoneal injection of 7.5 IU pregnant mare serum gonadotropin (ProSpec, HOR-272) followed by 7.5 IU human chorionic gonadotropin (HCG) (ProSpec, HOR-250) with a 48 hours interval. Immediately after the HCG injection, female mice were housed with DBA/2 male mice for mating. Zygotes were collected from the oviducts and treated with 0.1% hyaluronidase (Sigma, H35056) in an M2 medium (Sigma, M7167) to remove cumulus cells. They were then transferred to KSOM medium (Millipore, MR-106-D) with paraffin oil (Sigma, 18512) at 37 °C under a 5% CO_2_ environment. After injection of 100 pg dsRNAs with or without 50 pg eGFP mRNA into the cytoplasm using a constant flow system (IX73 OLYMPUS and FemtoJet Eppendorf), embryos were cultured in KSOM medium to stage E4.5. The mouse E4.5 ESC line derivation was previously described.^47^ They were cultured in 2i medium (Millipore ESG1107) supplemented with 15% FBS (GIBCO). Two sgRNAs targeting the *Prkra* locus were introduced into mESCs,^48^ and mutant cells were obtained by FACS sorting and expanded to establish stable cell lines. Mutation was confirmed by Sanger sequencing and western blot analysis.

#### mRNA and dsRNA synthesis

The coding sequences from genes of interest were PCR amplified or directly synthesized and subcloned into the pCS2 vector via a modified Gibson Ligation method.^49^ Recombinant vectors were linearized by NotI digestion, purified with a PCR cleanup kit, and used as the DNA templates for mRNA synthesis using the mMESSAGE mMACHINE SP6 or T7 kit (Cat. #AM1344 and Cat. #AM1340) according to the manufacturer’s instruction. The DNA template was removed by DNase I treatment at 37 °C for 15 minutes, and synthesized mRNAs were extracted by pheno-chloroform and precipitated by adding an equal volume of ice-cold isopropanol. After centrifugation at 12,000 *g* at 4 °C for 15 minutes, the pellet was rinsed with 70% cold ethanol and resuspended in 15-20 μL RNase-free water.

Templates for dsRNAs were PCR amplified from plasmids containing the entire ORFs or synthesized directly. T7 and SP6 promoter sequences were added to the 5’ or 3’ end of the DNA template by incorporating them into the 5’ end of amplifying primers. Sense and antisense RNAs were synthesized separately using T7 or SP6 RNA polymerase. They were annealed in a PCR thermocycler with a program of 95 °C for 5 min, ramping down at 0.1 °C/s to 50 °C and quickly cooling to 4 °C.

#### Microinjection

Microinjection was performed using a PLI-100A Picoliter microinjector (Harvard apparatus). A volume of 1-2 nL was injected into the embryo at the 1-cell stage.

#### Western blotting

Embryos were dechorionated, and the yolk was removed manually. Cell pellets were collected in 1.5 mL centrifuge tube and lysed in a lysis buffer (100 mM NaCl, 0.5% NP-40, 5 mM EDTA, 10 mM Tris-HCl, pH 7.5) containing protease inhibitors. Samples were boiled in Lammeli buffer (10% SDS, 0.5% bromophenol blue, 50% glycerol, 5% β-mercaptoethanol, 250 mM Tris-HCl, pH 6.8). Proteins were separated by SDS polyacrylamide gel electrophoresis and transferred to a nitrocellulose membrane. Primary antibodies used for western blot analysis are listed in the key resources Table.

#### qRT-PCR

Total RNAs were extracted at appropriate stages, and cDNAs were prepared using the TransScript First-Strand cDNA Synthesis Kit (Transgene) following the manufacturer’s protocols. qPCR was performed using 2 x M5 HiPer SYBR premix (MF787-01, Mei5) on a Q2000 real-time PCR system (LongGene). Primers used for amplifying genes of interest are listed in Supplementary Table S2. The 2^-ΔΔCt^ method was used to estimate relative expression levels with *gapdh* amplicon as the loading control.

#### ISH

ISH was performed as described.^50^ Templates for synthesizing *prkra* probes were amplified by PCR. Digoxigenin-labeled probes were *in vitro* transcribed using the DIG-labeling mix (Roche) in the presence of SP6 RNA polymerase. After DNase I treatment, labeled probes were directly precipitated with 50% v/v isopropanol.

#### Puromycin incorporation assay

Embryos were treated with 50 μg/mL puromycin from 2 hpf to 4 hpf. They were harvested following removal of yolk cells and lysed in protein IP buffer (10 mM Tris-HCl, pH 8, 150 mM NaCl, 1% NP-40, 1 mM EDTA, 10 μg/mL Aprotinin, 10 μg/mL Leupeptin, and 1 mM PMSF) After denaturation in 1x protein loading buffer at 95 °C for 5 minutes, the puromycin-conjugated proteins were visualized by western blot analysis using a puromycin antibody.

#### Luciferase assay

Synthetic *luciferase* mRNAs (50 pg) were injected alone or coinjected with dsRNA or dsRNA-contaminated mRNAs into 1-cell stage embryos. At 6 hpf, injected embryos (30 at least) were lysed in a passive lysis buffer provided in the Dual-Luciferase Reporter Gene Assay kit. After centrifugation at 12,000 *g* for 10 minutes, the supernatant was mixed with firefly luciferase substrate (50 μL lysate plus 50 μL substrate) and quantified using a luminometer (Berthold, Junior LB 9509).

#### Sucrose density gradient centrifugation

Sucrose was thoroughly dissolved in polysome buffer (50 mM Tris-HCl, pH 7.0, 100 mM NaCl, 5 mM MgCl_2_, 100 μg/mL CHX) to prepare 10% and 60% sucrose solutions. The sucrose density gradient was generated in a 41Ti centrifuge tube on a Gradient Station (BIOCOMP) and allowed to equilibrate. Embryos at 5-hpf were collected from controls and dsRNA-injected groups. After removal of the chorions using pronase, embryos were treated with 50 μg/mL CHX for 10 minutes and lysed on ice in lysis buffer (1% Triton X-100, 300 U/mL RNase inhibitor, 10 μg/mL aprotinin, 10 μg/mL leupeptin, and 1 mM PMSF). The lysate was centrifuged at 12,000 *g* for 10 minutes at 4 °C. The supernatant was carefully loaded onto the top of the sucrose density gradient curtain and centrifuged at 240,000 g for 2 hours using an ultracentrifuge (Beckman, XPN-100). Following centrifugation, the centrifuge tube was loaded onto a density fractionation system (BIOCOMP), which was utilized to separate the fractions and measure the absorbance at 260 nm. A 10 kDa ultrafiltration tube (Millipore) was used to concentrate proteins according to the manufacturer’s instructions.

#### Ribosome profiling and bioinformatic analysis

This was performed by following established experimental procedures with modifications.^51^ Briefly, both control and dsRNAs-injected embryos at 5 hpf were thoroughly lysed on ice in lysis buffer (20 mM Tris-HCl, pH 7.4, 150 mM NaCl, 5 mM MgCl_2_, 1 mM DTT, 100 µg/ml CHX, 1% Triton X-100 and 25 U/mL Turbo DNase I). The lysate was centrifuged at 12,000 *g* at 4 °C for 10 minutes to remove cell debris. The supernatant was added with 0.2 U/μL RNase I and incubated at 25 °C for 30 minutes to digest free RNAs unprotected by ribosomes. The digestion was terminated by adding 0.67 U/μL RNase inhibitor. The lysate was then subjected to sucrose density gradient centrifugation, and fractions with a single absorbance peak at 260 nm, corresponding to the 80S monosome, were collected and thoroughly lysed in TRIzol for RNA extraction. RNA fragments within the size of 26-34 nt were separated by denaturing polyacrylamide gel electrophoresis, and desired bands were recovered in RNA elution buffer (300 mM sodium acetate, pH 5.5, 1 mM EDTA, 0.25% SDS). Eluted RNAs were dephosphorylated by T4 PNK and subjected to ligation reaction using T4 RNA ligase 2, to add a preadenylylated and 3’-blocked linker (5’ rApp-CTGTAGGCACCATCAAT-NH_2_ 3’) at the 3’ end. The RNA fragments with linker were extracted and reverse transcribed using SuperScript III in the presence of the primer 5’ (Phos)-AGATCGGAAGAGCGTCGTGTAGGGAAAGAGTGTAGATCTCGGTGGTCGC-(SpC18)-CA CTCA-(SpC18)-TTCAGACGTGTGCTCTTCCGATCTATTGATGGTGCCTACAG-3’ to obtain extended cDNA molecules. Purified cDNAs were subjected to circularization using CircLigase, forming circular cDNA molecules. To remove residual rRNA components in the cDNAs, we adopted a method involving hybridization with biotinylated rRNA-depletion oligo primers specific to zebrafish rRNAs (GGGGGGATGCGTGCATTTATCAGATCA, TTGGTGACTCTAGATAACCTCGGGCCGATCGCACG, GAGCCGCCTGGATACCGCAGCTAGGAATAATGGAAT, TCGTGGGGGGCCCAAGTCCTTCTGATCGAGGCCC, GCACTCGCCGAATCCCGGGGCCGAGGGAGCGA, GGGGCCGGGCCGCCCCTCCCACGGCGCG, GGGGCCGGGCCACCCCTCCCACGGCGCG, CCCAGTGCGCCCCGGGCGTCGTCGCGCCGTCGGGTCCCGGG, TCCGCCGAGGGCGCACCACCGGCCCGTCTCGCC, AGGGGCTCTCGCTTCTGGCGCCAAGCGT, GAGCCTCGGTTGGCCCCGGATAGCCGGGTCCCCGT, GAGCCTCGGTTGGCCTCGGATAGCCGGTCCCCCGC, TCGCTGCGATCTATTGAAAGTCAGCCCTCGACACA, TCCTCCCGGGGCTACGCCTGTCTGAGCGTCGCT). The remaining circularized cDNAs were amplified through PCR using forward library primer (5’-AATGATACGGCGACCACCGAGATCTACAC-3’) and reverse indexed primer (5’- CAAGCAGAAGACGGCATACGAGATNNNNNNGTGACTGGAGTTCAGACGTGTGCTCTTCCG -3’, with NNNNNN representing the reverse complementary sequence of the barcode). The amplified products were separated and recovered using polyacrylamide gel electrophoresis with TE buffer. Subsequently, the library was subjected to quality control assessment, and Illumina Novaseq 6000 S4 (PE150) sequencing was performed to obtain the sequences of ribosome-protected RNA fragments.

Cutadapt was utilized for removing adapters and low-quality reads, Bowtie2 was employed for data mapping and Samtools was used to remove duplicated reads in mapped RNA-seq data.^52–54^ Self-written scripts were utilized to read and count Ribo-seq and RNA-seq mapping data, analyze RPF position and ribosome density, and visualize the results.

Transcriptome and annotation data were downloaded from NCBI (GCF_000002035.6_GRCz11_rna_from_genomic.fna, GCF_000002035.6_GRCz11_genomic.gtf). The non-coding RNA and mRNA sequence datasets are manually extracted from the downloaded FASTA files.

For Ribo-seq data, adapters and low-quality bases were removed from reads using the cutadapt arguments’ -a ACGACGCTCTTCCGATCT…CTGTAGGCACCATCAATAGATCGG -A ATTGATGGTGCCTACAG…AGATCGGAAGAGCGTCG -q 15,10’; reads lengths less than 20 nt or greater than 40 nt after adapter cutting were removed using cutadapt arguments ‘--minimum-length 20 --maximum-length 40’. For RNA-seq data, adapters, low-quality bases and Ns were removed from the reads using the cutadapt arguments’ -a AGATCGGAAGAGCACACG -A AGATCGGAAGAGCGTCGT -q 15,10 --trim-n’; reads length less than 50 nt after adapter cutting were removed using cutadapt argument ‘--minimum-length 50’.

The clean data of forward reads from Ribo-seq was mapped to non-coding RNA data using Bowtie2 with the arguments “--un unmapped. fasta -k 3”, and unmapped reads were subsequently mapped to mRNAs using the arguments “-k 3” to generate the SAM file. Reads mapped to multiple genes were assigned randomly, with probabilities proportional to the number of unique reads associated with each gene. Reads mapped to multiple gene IDs with the same gene name were assigned to the gene ID with the highest number of unique reads. Finally, the reads mapped to multiple transcripts with the same gene ID were assigned to the transcript with the highest number of unique reads.

The clean RNA-seq data were mapped to the transcriptome using Bowtie2 with the argument “-k 3”. Duplicated reads were removed from the mapped data using Samtools with the commands: ‘samtools fixmate -m; samtools sort; samtools markdup; samtools sort’. Read mapped to multiple gene names, gene IDs, and transcript IDs were handled using the same strategy described for mapping Ribo-seq reads.

The position of RPFs/read on the transcript was determined by the position of the 5’ end of the read. The positions of codons on the transcript were identified using the position of the first nt relative to the 5’ end of the transcript, with the first nt being designated as position 1. The relative position of a read to the start or stop codon was calculated by subtracting the position of the start or stop codon from the position of the read, respectively. The number of reads relative to the start codon ranging from -24 to 24 nt was counted. In addition, the number of reads was counted relative to the stop codon, ranging from -36 to 12 nt. The number of reads was visualized in a bar plot, and the frame of a read was determined by the remainder by dividing its relative position to the start codon by 3. Subsequently, the fraction of reads, with relative positions to start codon ranging from -12 to 72 nt, that fall into different frames was calculated and visualized in another bar plot.

Reads with relative positions to the start codon between -100 and 300 nt and relative positions to the stop codon between -300 and 100 nt were counted. Only transcripts with CDSs longer than 600 nt were considered during the analysis. A distribution of the number of reads along the mRNA was plotted.

For ribosome density analysis, the following formula is employed:^35^

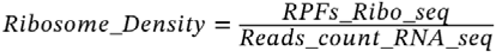

The sum of RPF/Read count data was scaled to 1e6 to mitigate the effects of sequencing depth. The ribosome density was calculated for genes only with a read count exceeding 64.

Given that the injection of dsRNAs alters the overall translational activity, as demonstrated by the puromycin incorporation experiments, a correction factor was introduced to accommodate this alteration (0.5 for 30 pg/embryo dsRNA injection; 0.3 for 100 pg/embryo injection). The ribosome density of the dsRNA-injected group was multiplied by this factor to reflect the overall translational change in the treatment. Fold changes in ribosome number (the RPF number) after dsRNA stimulation indicate protein synthesis rate. The ribosome number of the treatment group was also multiplied by the correction factor.

GO analysis was performed using tools provided in DAVID Bioinformatics (https://david.ncifcrf.gov/tools.jsp),^55^ using all genes with RPF read counts of more than 64 as the background.

#### RNA pulldown

A detailed protocol is provided in the manual of the Pierce^TM^ Magnetic RNA-Protein Pulldown kit. In general, 50 μL of streptavidin magnetic beads were transferred to RNase-free centrifuge tubes. The tubes were placed on a magnetic separation stand for 30 seconds, and the supernatant was removed. The beads were washed twice with 20 mM Tris-HCl, pH 7.5 (50 μL for each sample). After adding 1x RNA Capture Buffer, the beads were vortexed and incubated with 50 pmol of biotin-labeled RNAs (for specific sequences, see Table S1) under gentle rotation at room temperature for 20 minutes. The supernatant was removed, and the beads were washed twice with 20 mM Tris-HCl, pH 7.5. In parallel, 400 de-yolked embryos were lysed in 400 μL protein lysis buffer (10 mM Tris-HCl, pH 8, 150 mM NaCl, 1% NP-40, 1 mM EDTA, 10 μg/mL aprotinin, 10 μg/mL leupeptin, and 1 mM PMSF). After centrifugation at 12,000 *g* for 10 minutes at 4 °C, the 130 μL cleared supernatant was diluted to a 400 μL system containing 1 x protein-RNA binding buffer and 15% glycerol, which was then incubated with the beads at 4 °C under rotation for 1 hour. Following this incubation, the beads were washed twice with 100 μL 1 x washing buffer with gentle mixing. Finally, the beads were incubated with 50 μL of elution buffer at 37 °C for 20 minutes. The eluent was subjected to mass spectrum and western blot analyses.

In specific experiments for Figures 3C and E, 4 μg recominant Prkra-6xHis were added to 400 μL diluted cell lysate and incubated at 28.5 C for 1 hour. The other RNA-pulldown procedure was carried out as described in the previous paragraph.

#### Mass Spectrometry

After RNA pulldown, the samples were subjected to polyacrylamide gel electrophoresis to separate the proteins. To visualize proteins, the gel was stained using a silver staining kit (Beyotime, Cat. #P0017S). The silver-stained proteins in each lane were then excised into four parts as independent samples with different molecular weights. Protein digestion was carried out using trypsin following the filter-aided sample preparation (FASP). Digested peptides obtained from each sample were desalted using 3 mL of Empore™ SPE Cartridges C18 (standard density, bed I.D. 7 mm; Sigma-Aldrich). Subsequently, they were concentrated using vacuum centrifugation and reconstituted in 40 µL of 0.1% (v/v) formic acid. LC-MS/MS analysis was conducted using a timsTOF Pro mass spectrometer (Bruker) coupled to Nanoelute (Bruker). The peptides were loaded onto a C18 reversed-phase analytical column (Thermo Scientific Easy Column, 25 cm long, 75 μm inner diameter, 1.9 μm resin) with 95% buffer A (0.1% formic acid in water) and separated using a linear gradient of buffer B (99.9% acetonitrile and 0.1% formic acid) at a flow rate of 300 nL/minute. The mass spectrometer operated in positive ion mode with an applied electrospray voltage of 1.5 kV. Precursor ions and fragments were analyzed using the TOF detector within a mass range of m/z 100-1700. The timsTOF Pro operated in parallel accumulation serial fragmentation (PASEF) mode, with PASEF data collection performed under the following parameters: ion mobility coefficient (1/K0) ranging from 0.6 to 1.6 Vs cm2; 1 MS scan and 10 MS/MS PASEF scans were acquired. Active exclusion was enabled with a release time of 24 seconds.

Two negative controls (RNA-free beads and ssRNAs) were included to set up a stringent strategy for screening dsRNA-associated proteins. For specific sequences of ssRNA and dsRNA used for RNA-pulldown/MS, see Table S1. The criteria for identifying a specific dsRNA-associated protein are as follows: relative intensity (RI) for dsRNA pulldown (RI_dsRNA_) should be greater than 10e-4, the ratio of RI for dsRNA (RI_dsRNA_) to RI for ssRNA (RI_ssRNA_) should be greater than 3, and the ratio of RI for beads (RI_beads_) to the average RI of dsRNA and ssRNA ((RI_dsRNA_+RI_ssRNA_)/2) should be less than 0.2. Here, RI is defined as the total intensity of a specific protein divided by the sum of the intensity of all detected proteins. Scatter plot in Figure 2B was generated with the horizontal axis representing RI_dsRNA_, while the vertical axis representing the enrichment index defined as log_10_(1/(RI_ssRNA_/RI_dsRNA_+0.01)). Specific dsRNA-associated proteins were marked in red, whereas grey dots designate proteins with RI_dsRNA_/RI_ssRNA_≦3 and RI_beads_/((RI_dsRNA_+RI_ssRNA_)/2)<0.2, and grey circles stand for proteins with RI_beads_/((RI_dsRNA_+RI_ssRNA_)/2)≧0.2.

#### Recombinant protein expression and purification

The pET-Duet plasmid was linearized using KpnI and NcoI. A 6 x His tag sequence was inserted at the 3’ end in the coding regions of *prkra* or its mutants. The *Rossta* strain of *E. coli* transformed with plasmids was cultivated overnight at 37 °C in LB and inoculated to 1 liter of fresh LB for amplification until reaching 0.6 OD (optical density). To induce protein expression, IPTG (1 mM, final concentration) was added to the culture for another 26 hours at 20 °C with agitation. The bacteria were then harvested by centrifugation at 2,800 g for 10 minutes at 4 °C, and 10% glycerol was used to resuspend the pellets, followed by centrifugation at 1,850 g for 20 minutes at 4 °C. Cell pellets were weighed and resuspended in AP buffer (50 mM Tris-HCl, pH 7.2, 400 mM NaCl, 40 mM imidazole, 10% glycerol, and 0.02% NP-40). Cells were homogenized using lysozyme solution (1 mM DTT, 1 mM PMSF, and 1 mg/mL lysozyme in AP buffer), and sonication was performed using an ultrasonic homogenizer (JY92-IIN) with the program: 5 seconds ON and 10 seconds OFF cycle for 4 minutes, with power setting 20%. After centrifugation at 24,000 *g* at 4 °C for 1 hour, the supernatant was filtered using a 0.45 μm membrane. In parallel, the Ni column was prepared and washed twice with ddH_2_O, followed by one wash with AP buffer. The cleared supernatant was added to the column and incubated at 4 °C for 1 hour to allow protein binding. The column was then drained and washed twice with buffer A (1 mM DTT and 1 mM PMSF in AP buffer), followed by the addition of cold buffer B (2 mM ATP-Na, 1 mM MgAc, 1 mM DTT, and 1 mM PMSF in AP buffer) and incubation at 4 °C for 10 minutes to activate the protein. After two washes with buffer C (1 mM DTT in AP buffer), proteins were slowly eluted with the elution buffer (50 mM Tris-HCl, pH 7.2, 400 mM NaCl, 400 mM imidazole, 10% glycerol, and 0.02% NP-40). For long-term storage, glycerol was added to the proteins at a final concentration of 50% and stored at -80 °C.

#### *In vitro* translation

*In vitro* translation was conducted using the Promega reticulocyte *in vitro* translation system (catalog number L4960), by following the manufacturer’s instructions. A total of 300 ng of luciferase mRNA, with or without the addition of 300 ng dsRNAs, were incorporated into a 12 μL reaction mixture. This mixture was then incubated at 30 °C for 90 minutes. To assess the impact on translation efficiency, either wild-type Prkra-6xHis or the Δ123-6xHis mutant proteins were added to the *in vitro* translation system at a dose of 1 μg. The luciferase activity was analyzed as described.

#### EMSA assay

Biotin-labeled 139 bp RNA was from *Vpr* gene of HIV-1 with the sequence: UCCUAGGAUUUGGCUCCAUGGCUUAGGGCAACAUAUCUAUGAAACUUAUGGGGAUACUUGGGCAGGAGUGGAAGCCAUAAUAAGAAUUCUGCAACAACUGCUGUUUAUCCAUUUCAGAAUUGGGUGUCGACAUAGCAGA. Forward RNA strands were *in vitro* transcribed with biotin-labelling mix (Roche, 75501920) and annealed with unlabeled reverse strands. The dsRNAs and protein were mixed in 1 x binding buffer (10 x storage: 80% glycerol, 100 mM MgCl_2_, 5 M NaCl, 0.5 M EDTA, 1 M DTT, 1 M Tris-HCl, pH 8.0), followed by incubation at room temperature for 15 minutes. Subsequently, the protein-RNA mixture was electrophoresed using a 6% native-PAGE gel at 80 V for 2 hours in 1x running buffer (25 mM Tris, 0.5 M glycine) and then transferred onto a nylon membrane using a transfer apparatus and 0.5 x TBE buffer for 4 hours. UV cross-linking was performed at a dose of 0.12 J/cm² using a 254 nm UV source for 50 seconds to immobilize the complex on the nylon membrane. Finally, biotin-labeled RNAs were detected using stabilized streptavidin-HRP.

#### Blue native gel electrophoresis

A 4-13% Blue Native gel (Beyotime, P0546S) was assembled in the Mini-PROTEAN System (BIO-RAD). The inner and outer chambers were filled with BN cathode buffer I (Beyotime, P0765S) and BN anode buffer (Beyotime, P0769S), respectively. Pre-electrophoresis was then performed at 100 V for 1 hour. During this time, recombinant Prkra-6xHis was incubated in Protein-RNA binding buffer (10× storage: 80% glycerol, 100 mM MgCl_2_, 5 M NaCl, 0.5 M EDTA, 1 M DTT, 1 M Tris-HCl, pH 8.0) for 15 minutes at room temperature. The mixture was then gently combined with Coomassie Brilliant Blue sample buffer (Beyotime, P0761S) before being loaded onto the BN gel and electrophoresed for 15 minutes. At this point, the cathode buffer I in the inner chamber was replaced with cathode buffer II (Beyotime, P0767S), and electrophoresis continued for an additional 30 minutes. The BN gel was subsequently removed from the system and rinsed overnight in 10% acetic acid on a horizontal shaking table. Finally, gel imaging was performed using a full-spectrum gel imager (Invitrogen, iBright 1500).

#### RNA-IP

Dyna Protein G beads (25 μL) were added to each sample in an RNase-free microcentrifuge tube, which was placed on a magnetic stand for 30 seconds. After careful removal of the supernatant, the beads were washed twice with 1 mL of IP-blocking buffer (0.5% BSA in PBS) and incubated with 25 μL of IP-blocking buffer and 3 μL anti-His antibody overnight at 4 °C with rotation to facilitate antibody binding. The antibody-coated beads were then washed thrice with the IP-blocking buffer.

In parallel, the Prkra-His protein and biotin-labeled dsRNAs were incubated in a protein-RNA binding buffer at room temperature for 20 minutes in a 50 μL reaction volume. After adding 500 μL of 1 x binding buffer and 25 μL of prepared anti-His beads, the mixture was incubated at room temperature under rotation for 30 minutes. Subsequently, the magnetic beads were gently washed thrice with 1 x binding buffer, followed by adding TE buffer. The grasped biotin-labeled RNAs were released by boiling at 95 °C for 5 minutes. Dot blot analysis was performed using the stabilized streptavidin-HRP conjugate to visualize Prkra interacting RNAs.

#### Co-IP

Anti-His magnetic beads (Biolinkedin, L-1015) were used for immunoprecipitation experiments. They were washed thrice and stored at 4 °C in the wash buffer (20 mM Tris-HCl, pH 7.5) before use. For Figure 3E, synthetic mRNAs (100 pg) were injected into 1-cell stage embryos, and approximately 150 injected embryos were collected around 8 hpf, followed by dechorionation and deyolking as previously described.^56^ After adding 200 μL of lysis buffer containing 10 mM Tris-HCl, pH 7.5, 150 mM NaCl, 0.5 mM EDTA, 0.5% NP-40, and 1x protease inhibitor cocktail (Roche, 40609), the embryos were homogenized using a grinding stick on ice for 10 minutes, followed by centrifugation at 12,000 g for 10 minutes at 4 °C. After saving 20 µL as input control, the supernatant was incubated with 10 µL of beads at 4 °C overnight. The beads were washed three times with lysis buffer, boiled at 95 °C for 5 minutes, and subjected to western blot analysis.

For Co-IP experiments in Figure 4D and E, 4 μg of recombinate Prkra-6xHis with or without 1 μg dsRNAs were added to 200 μL cell lysate made from 200 embryos at 4 hpf in lysis buffer containing 2 x protease inhibitor cocktail (Roche, 40609), and incubated for 2 hours at 28.5 °C. And then the immunoprecipitation experiments were performed as described above.

#### Toeprinting assay

This was performed by following established experimental procedures with modifications.^57,58^ Briefly, this experiment comprises three steps: preparation of lysate for *in vitro* translation, *in vitro* translation coupled with reverse transcription, and visualization of toeprinting ssDNA products.

For cell lysate preparation, embryos at 4 hpf were dechorionated, and the yolk was manually removed. Cell pellets were collected in Eppendorf tubes and suspended in an equal volume of cold lysolecithin lysis buffer (20 mM HEPES, pH 7.5, 100 mM potassium acetate, 1 mM magnesium acetate, 2 mM dithiothreitol and 100 mg/mL L-α-lysophosphatidylcholine from Merck, Y0001904) for 1 minute. The embryonic cells were then pelleted by centrifugation at 300 *g* for 30 seconds. Subsequently, the pellets were resuspended in an equal volume of ice-cold hypotonic extraction buffer (20 mM HEPES, pH 7.5, 10 mM potassium acetate, 1 mM magnesium acetate, 4 mM dithiothreitol, and Complete Protease Inhibitor) and incubated for 5 minutes. The cells were then thoroughly homogenized by repeated pipetting with a 20 μL pipette tip. Following this, the cell lysate was cleared by centrifugation at 10,000 *g* for 5 minutes at 4 °C. The resulting supernatant was prepared for *in vitro* translation and could also be stored in aliquots at -80 °C.

For *in vitro* translation and reverse transcription, 500 ng/μL of *gfp* mRNA was hybridized with 200 ng/μL of FAM-labeled primer (5’FAM-GATGAACTTCAGGGTCAGCTTG-3’) and mixed in equal volumes. The mixture was then incubated in a thermocycler using a hybridization program consisting of an initial step at 95 °C for 2 minutes, followed by slow cooling to room temperature. The translation reactions were performed in a 16 μL mixture containing 6 μL of cell lysate, 2 μL of the annealed mRNA/primer complex, and 8 μL of 2 x translation buffer (40 mM HEPES-KOH, pH 7.6, 160 mM potassium acetate, 2 mM magnesium acetate, 2 mM ATP, 0.24 mM GTP, 34 mM creatine phosphate from Sigma, CRPHO-RO, 0.2 mg/mL creatine phosphokinase from Shyuanye, S10076, 4 mM dithiothreitol, 40 mM of each of the 20 amino acids, 0.3 mM spermidine from Sigma, S2626, 200 μg/mL cycloheximide, and 800 U/mL RNase inhibitor). After thorough mixing, the reaction was incubated at 37 °C for 2 hours. The reverse transcription reaction was initiated by adding 6 μL of 5 x RT buffer (5 x PBS, 15 mM MgCl_2_), 1 μL of SuperScript III (Invitrogen, Cat. #18080-093), and 1 μL of dNTPs (10 mM each). The mixture was incubated at 37 °C for an additional hour. Markers for run-off and PIC were reverse transcribed using the same primer hybridized with either full-length or 5’ truncated *gfp* mRNA templates, generating cDNAs of 186 nt and 128 nt, respectively. The primers used for constructing vectors to generate RNA templates are listed in Table S2.

To visualize toeprinting products and markers, cDNA was extracted via precipitation with 50% isopropanol and washed twice with 70% ethanol. Following digestion with RNase H (Thermo, EN0202) for 30 minutes at 37 °C, the samples were mixed with a 2x loading buffer (98% formamide, 10 mM EDTA, 300 μg/mL bromophenol blue) and analyzed on a urea-PAGE gel (4% acrylamide/bisacrylamide 29:1, 8 M urea in 1 x TBE), using 1 x TBE as the electrophoresis buffer. The fluorescent bands in the gel were directly captured using a full-spectrum gel imager (Invitrogen, iBright 1500).

#### SVCV preparation and injection

Spring viremia of carp virus (SVCV) viral suspension stored at -80 °C was thawed on ice. Epithelioma papulosum cyprini (EPC) cells with a density of 80%-90% in serum-free M199 medium (Hyclone) were rinsed in cell culture flasks. The viral suspension was inoculated into the cells at a 1% volume ratio. The infection was carried out at 25 °C for 2 hours with intermittent shaking for 3-5 times. After viral infection, the viral fluid was collected, and the cell maintenance medium containing 2% FBS (Gibco) in M199 cell culture medium was added to the flasks for further cultivation. EPC cell infection was examined using an inverted microscope at 48-96 hours of cultivation. When more than 80% of cells exhibited cytopathic effects (CPEs), the flasks were subjected to three cycles of freeze-thawing in a -80 °C freezer, followed by centrifugation at 4 °C, 845 g for 10 minutes. The collected supernatant was aliquoted into 2 mL cryovials and stored at -80 °C for future use. EPC cells were seeded in a 96-well cell culture plate at a density of 1 x 10^4^ cells per well and cultured at 25 °C for 16-24 hours. When cell confluence reached 80%-90%, 100 μL of virus suspension at dilutions ranging from 10^-1^ to 10^-10^ was added to each well, with four replicates per dilution and three parallel experimental groups. The CPEs of monolayer cells at each dilution were examined every 24 hours. The semi-logarithmic Reed-Muench method was employed to calculate the half-tissue culture infectious dose (TCID_50_) of SVCV.^59^ Zebrafish embryos were injected with 2 nL of SVCV at a 5 x 10^6^ TCID_50_/mL concentration. Embryos were collected at 2-hour intervals, and the total viral load was quantified using qRT-PCR (primers in Supplementary Table S2). Subsequently, a viral growth curve was generated based on the levels of SVCV replication.

#### Protein structure prediction by Alphafold3

The online AlphaFold Server powered by AlphaFold 3 (https://golgi.sandbox.google.com/) was utilized to analyze the potential structures of the Prkra dimer and the Prkra-dsRNA complexes, following the guidelines provided in the online manual. The zebrafish Prkra protein sequence, with or without complementary 139 nt ssRNA sequences (see Table S1), was uploaded to the platform, and the number of Prkra units was set to either 2 or 8. The model with the highest confidence score was chosen for presentation and additional analysis. The dsRNA binding domains were subsequently highlighted using distinct colors in PyMOL Version 2.5.0a0.

#### Quantification and Statistical Analysis

All experiments were repeated at least two times with similar results. Data plotting and statistical tests were performed using GraphPad Prism 9. Statistical information is described in the figure legends.

