## Supplementary material for "Prkra dimer senses double-stranded RNAs to dictate global translation efficiency": Uncropped Blots

### Uncropped blots for Figure 1B

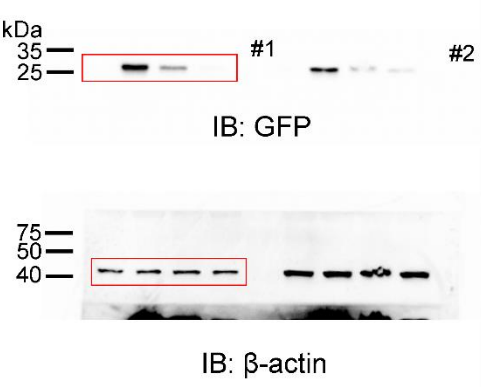


### Uncropped blots for Figure 1D

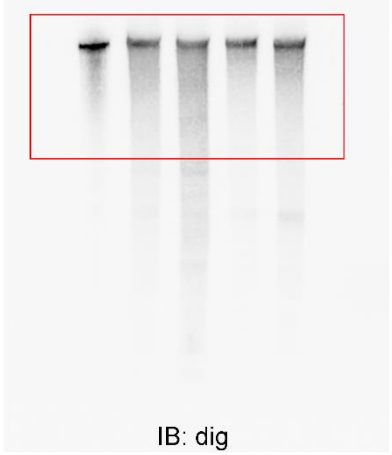


### Uncropped blots for Figure 1E

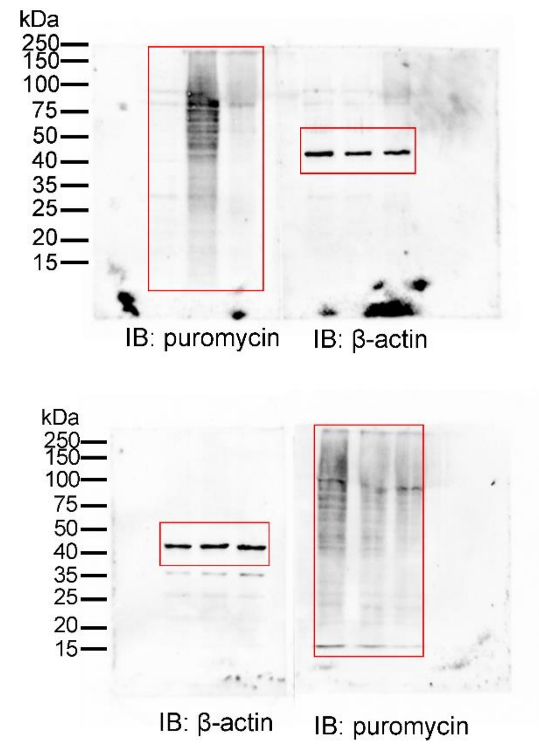


### Uncropped blots for Figure 1I

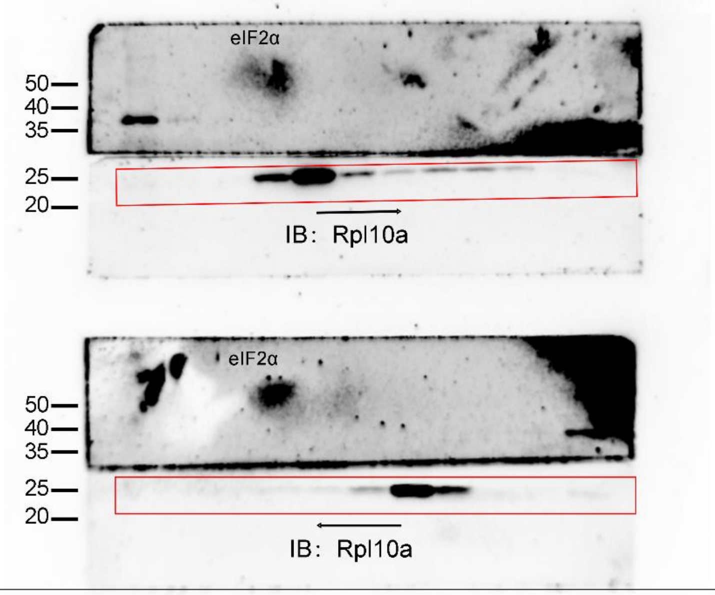


### Uncropped blots for Figure 1P

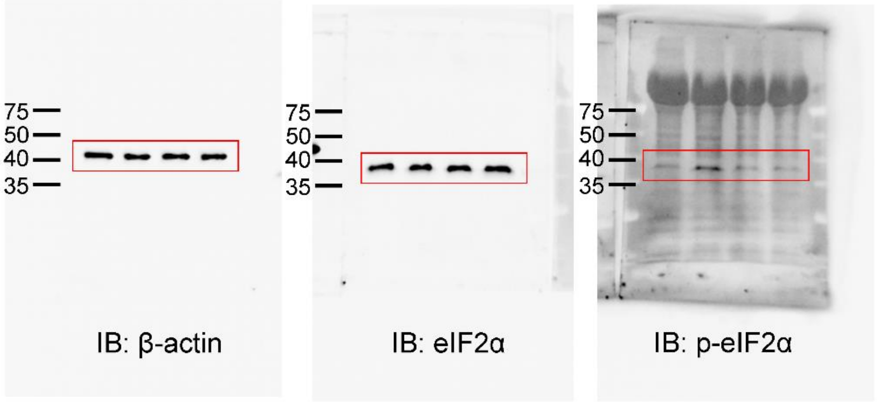


### Uncropped blots for Figure 1N

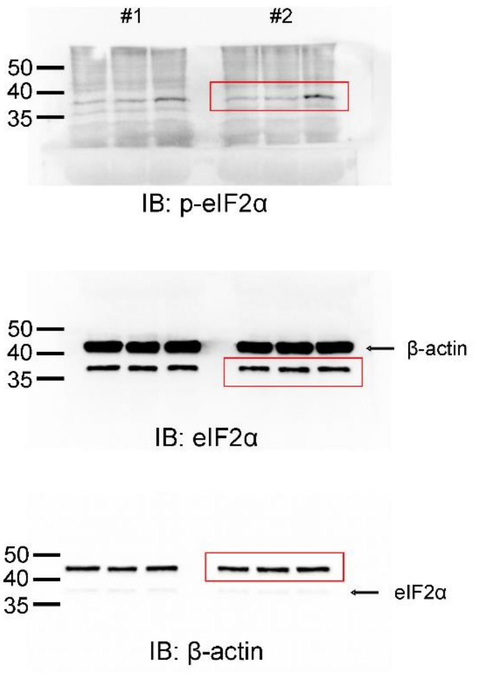

### Uncropped blots for Figure 2E

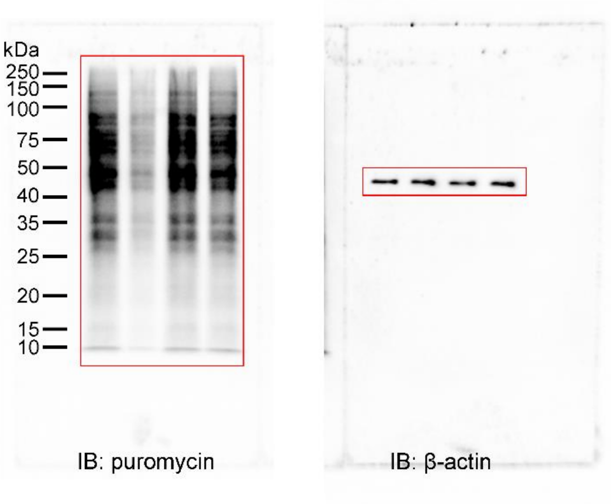


### Uncropped blots for Figure 3A

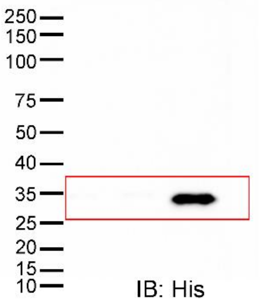


### Uncropped blots for Figure 3B

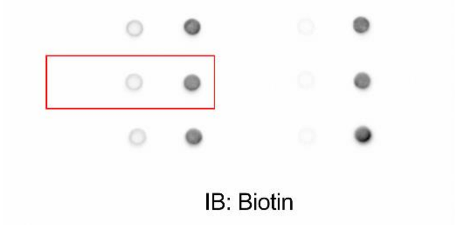


### Uncropped blots for Figure 3C

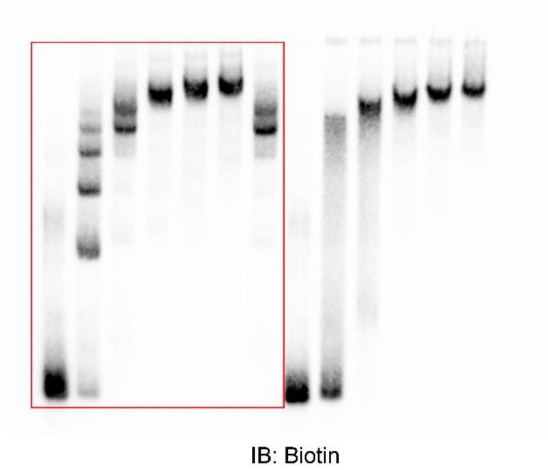


### Uncropped blots for Figure 3D

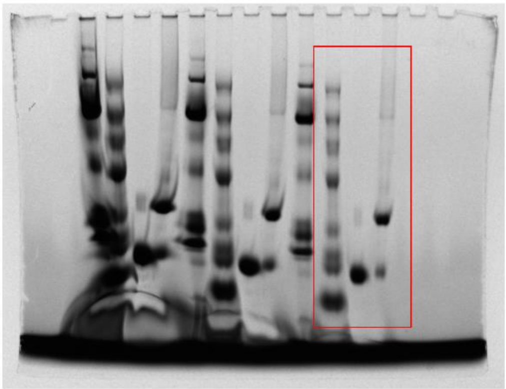

### Uncropped blots for Figure 3E

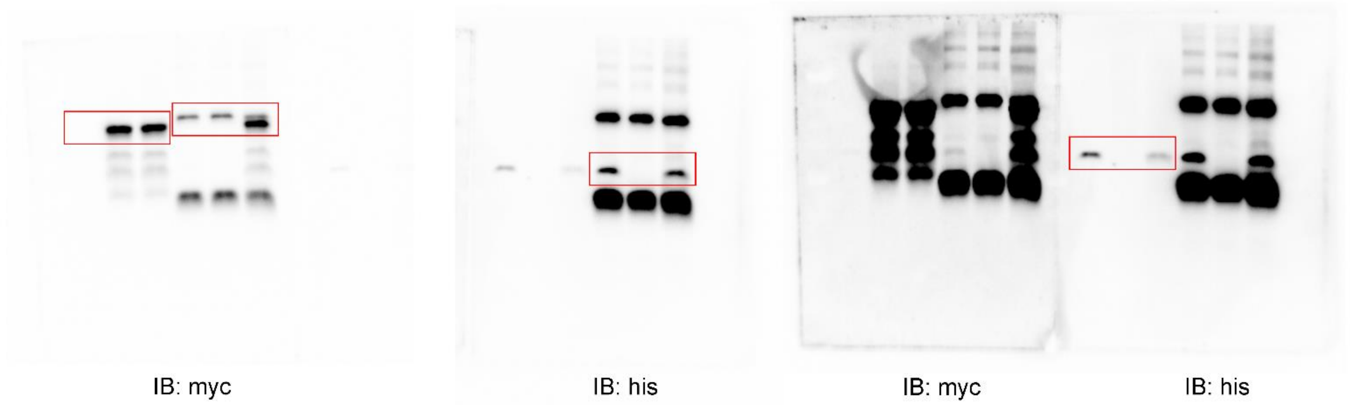

### Uncropped blots for Figure 3I

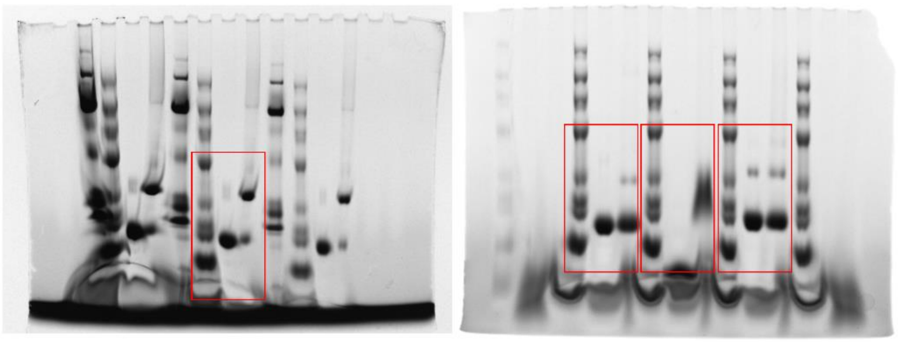

### Uncropped blots for Figure 3J

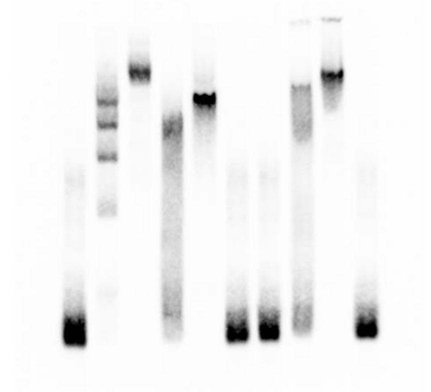

### Uncropped blots for Figure 4A

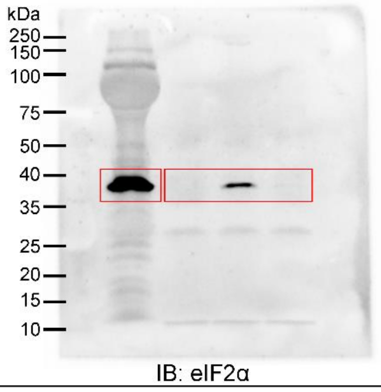


### Uncropped blots for Figure 4C

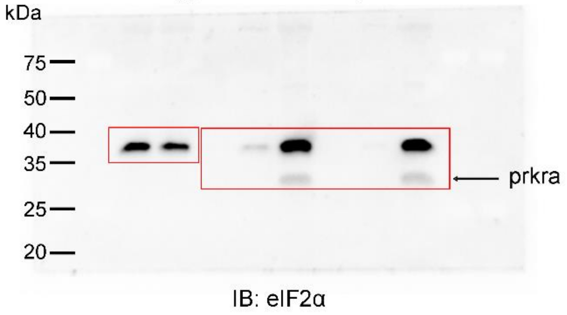


### Uncropped blots for Figure 4B

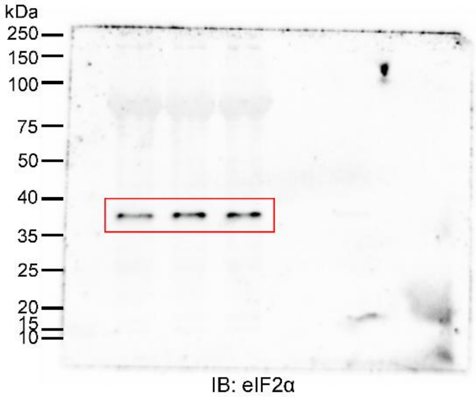


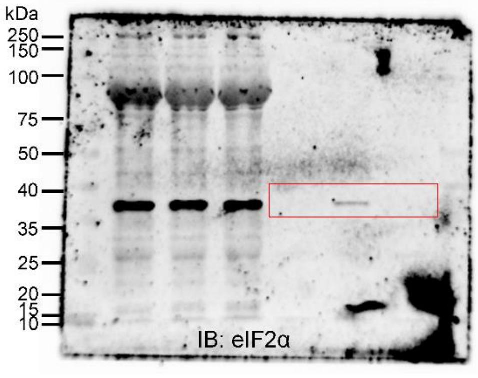


### Uncropped blots for Figure 4D

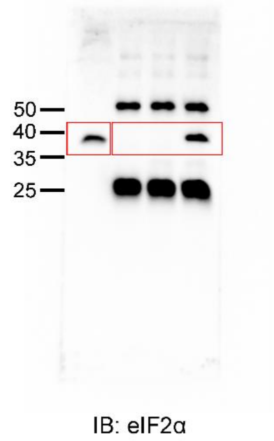


### Uncropped blots for Figure 4E

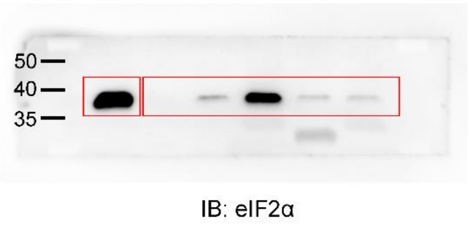


### Uncropped blots for Figure 4F

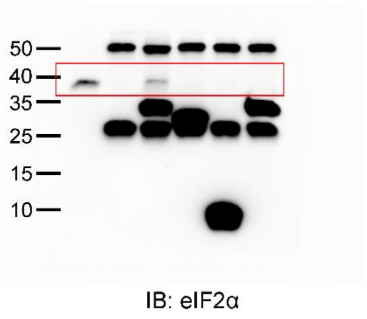


### Uncropped blots for Figure 4I

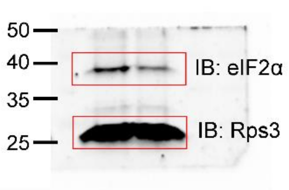

Uncropped blots for Figure 4K

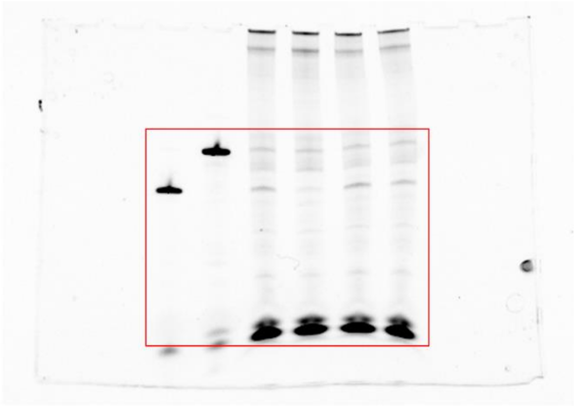

Uncropped blots for Figure 5D

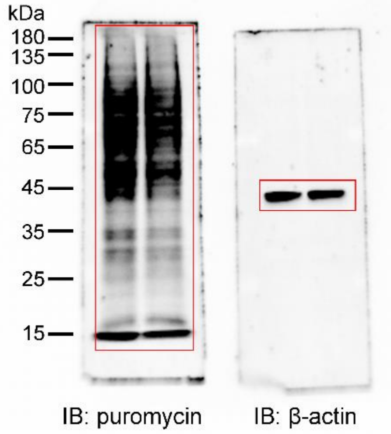


Uncropped blots for Figure 6B

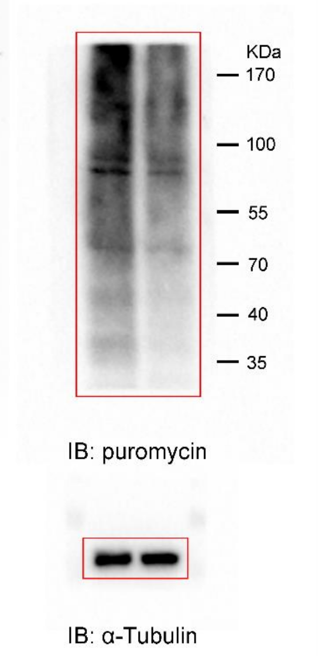


Uncropped blots for Figure 6F

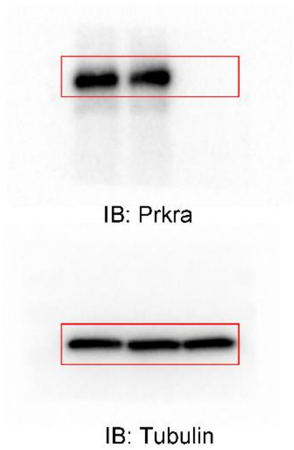


Uncropped blots for Figure 6H

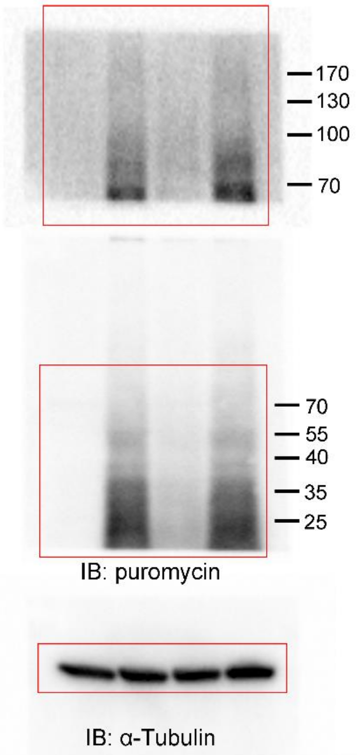


Uncropped blots for Figure S2F

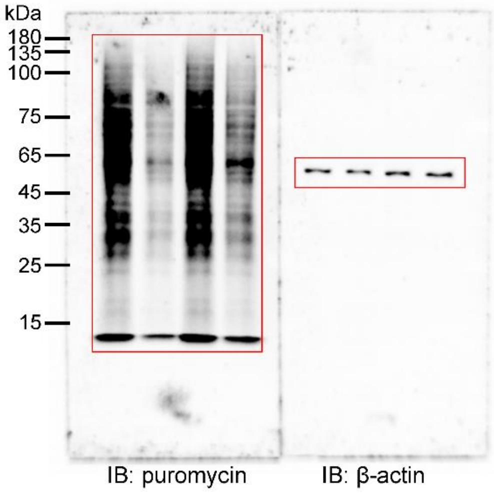


Uncropped blots for Figure S6

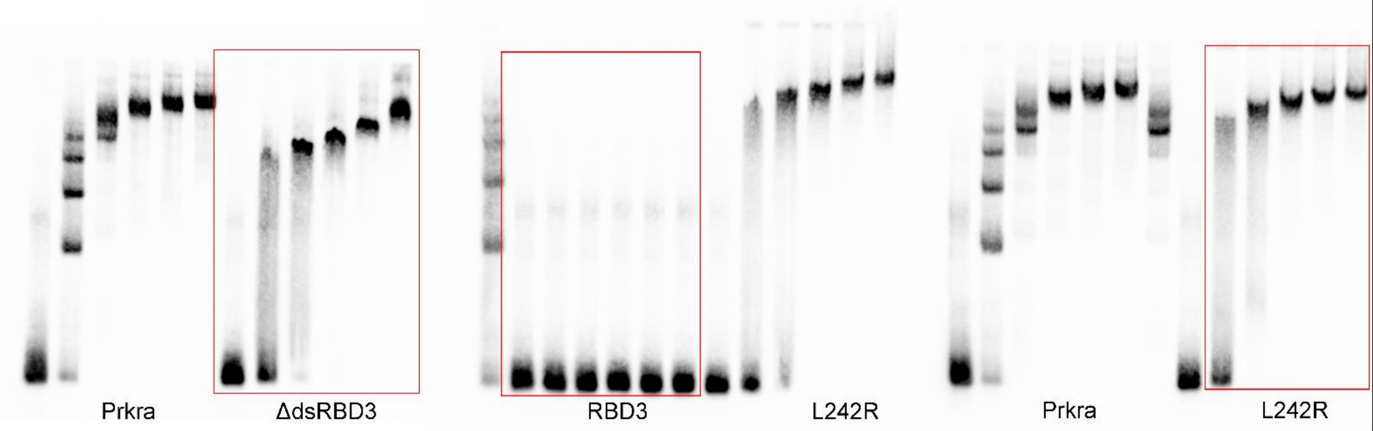
